# Neural transformation from image realism to perceived realness

**DOI:** 10.64898/2026.10.28.762323

**Authors:** Taiki Orima, Hiroshi Ban

**Author notes:** Correspondence: Taiki Orima.

## Abstract

Humans readily perceive whether a visual stimulus depicts something that could exist in the real world, yet the neural basis of perceived realness remains unclear. Because differences between real and unreal stimuli are ultimately expressed through visual features, realness might be expected to emerge along the ventral visual stream. Contrary to this prediction, we show that perceived-realness representations emerge in higher dorsal visual areas through a transformation of image-realism-related image-feature representations in early visual and early dorsal visual areas. fMRI revealed this representational shift toward intraparietal sulcus (IPS), with IPS patterns predicting perceived realness beyond image features and decision difficulty; source-localized MEG showed that realness coding in IPS peaked later than feature coding in early visual and dorsal visual areas. Together, these findings reveal a transformation from visual features to perceived realness along the dorsal visual stream. This representation may help guide subsequent actions and interactions with the real world.

## Introduction

Our visual system can easily perceive the extent to which a visual stimulus captures the real world. For example, a photograph captures the real world, whereas a realistic painting of the same scene can still be distinguished from the photograph, even when its composition and content are closely matched. Similarly, computer-generated (CG) images that faithfully follow the physical laws of the real world^1,2^ , as well as AI-generated images that have rapidly advanced in recent years, can appear highly realistic at first glance^3,4^, but they can sometimes evoke a sense of incongruity^4^. Such perceptual incongruity arising from highly realistic but imperfect artificial stimuli has also long been discussed in the context of the uncanny valley^5,6^. Thus, the perceived realness of a visual stimulus—that is, the extent to which the stimulus is perceived as capturing the real world—is a fundamental and important dimension of everyday visual perception. However, it remains unclear what neural mechanisms underlie this perception and how it arises.

How is perceived realness encoded in the human brain? The differences between real and unreal images ultimately arise from differences in pixel values and can be characterized in terms of image features derived from those pixel values. We therefore hypothesized that relatively low-level image features, such as image statistics, are encoded in the early visual system at early latencies^7–10^, followed by the representation of higher-level image features in higher-order visual areas, which are then transformed into a representation of perceived realness. Given a large number of previous studies showing the processing of image statistics, material information, and objects in ventral visual areas^11–13^, including the LOC^10,14–21^, it seems natural to suppose that the physical differences between real and unreal images—that is, their image features—are hierarchically encoded along the ventral visual stream and then transformed into a representation of perceived realness at later stages of that stream (Fig. 1a). However, the neural representations supporting object categorization may not directly account for perceptual distinctions between natural and synthetic images. For example, a previous study showed that observer models based on category-selective cortical responses and dCNN features could account for human object-category discrimination but failed to match human performance in distinguishing natural object images from synthetic images containing similar complex visual features in spatially scrambled arrangements^22^. Therefore, although ventral visual areas are plausible candidates, it remains unclear whether the transformation from image features to perceived realness is confined to the ventral visual stream. We therefore investigated the neural representation of perceived realness across a broad range of early, ventral, and dorsal visual areas. As a result, we found that neural representations associated with perceived realness emerged along the dorsal visual hierarchy and were particularly prominent in higher dorsal visual areas.

**Figure 1.**
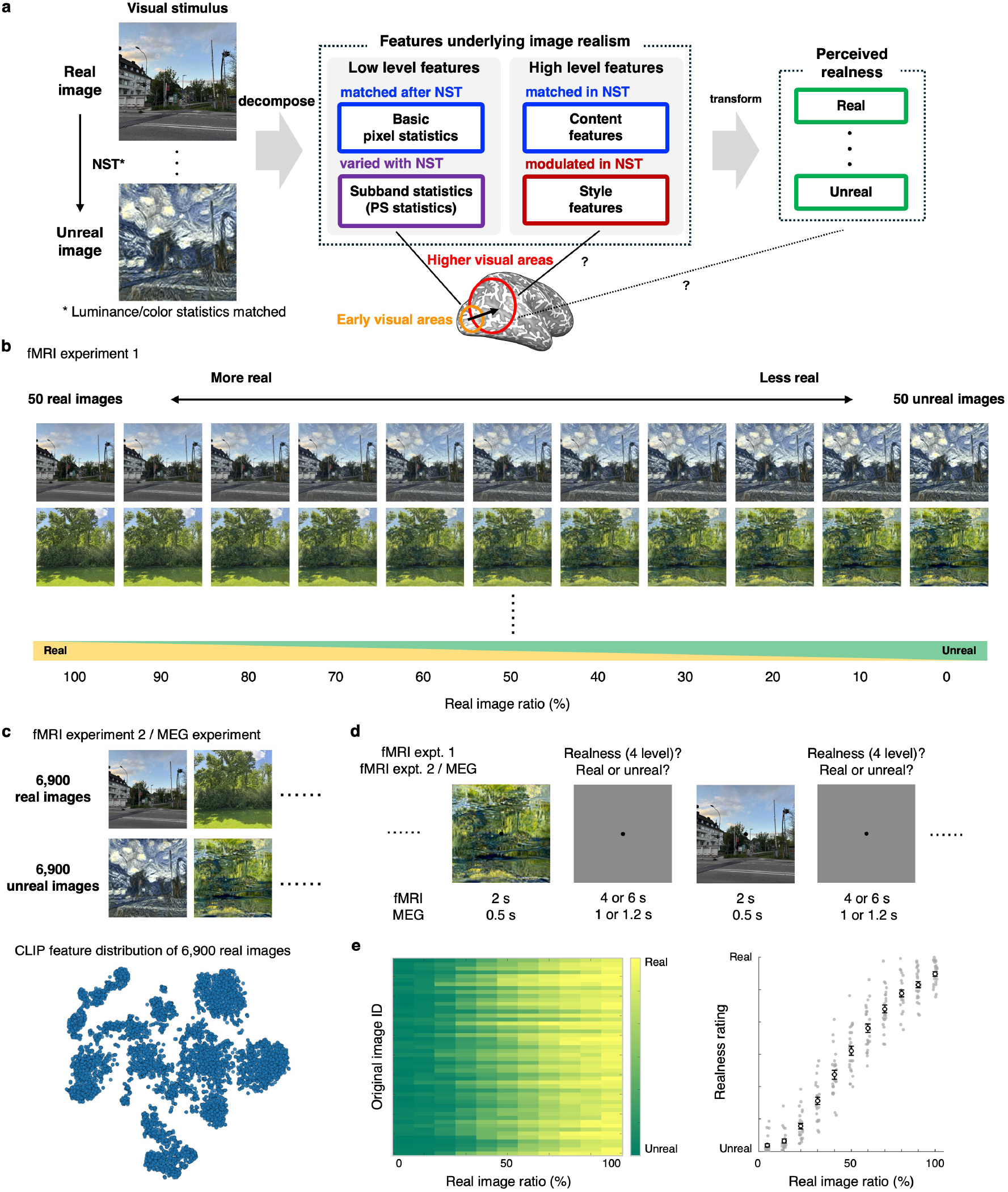
Hypothesis and visual stimuli. **(a)** We hypothesized that relatively low-level image features are processed in early visual areas, whereas higher-level image features are processed in higher visual areas, and that these representations are subsequently transformed into a representation of perceived realness. Based on this hypothesis, we investigated visual areas ranging from early visual areas to higher ventral and higher dorsal visual areas. **(b)** Visual stimuli used in fMRI experiments 1–1 and 1–2. We selected 50 real images from MS COCO^70^ and synthesized corresponding unreal images using neural style transfer^23^, with paintings selected from the refined WikiArt dataset^71^ as style images. We additionally generated intermediate images by linearly combining each real image with its corresponding unreal image at the pixel level. This procedure yielded 550 visual stimuli for the fMRI experiments. **(c)** Visual stimuli used in fMRI experiment 2 and the MEG experiment. A total of 6,900 real images were selected from MS COCO^70^, and 6,900 corresponding unreal images were synthesized using neural style transfer^23^, with paintings selected from the refined WikiArt dataset^71^ as style images. The t-SNE plot shows the CLIP features of the 6,900 real images and illustrates the diversity of the stimulus set in CLIP feature space. **(d)** Overview of the fMRI and MEG experimental procedures. In both experiments, a visual stimulus was presented at the center of the screen, followed by a uniform gray background during which participants made their responses. In the fMRI experiments, visual stimuli were presented for 2 s, followed by 4 or 6 s of gray background. In the MEG experiment, visual stimuli were presented for 500 ms, followed by 1,000 or 1,200 ms of gray background. **(e)** Behavioral results for perceived realness. The heatmap on the left shows the mean rating across participants for each stimulus. The plot on the right shows the mean ratings across participants and stimuli sharing the same real-image proportion. Both panels show that perceived realness changed systematically with the proportion of the real image.

The greatest challenge in investigating the neural representation of perceived realness is to construct a stimulus space in which image realism, or the visual fidelity of an image to the real world, can be varied continuously and systematically while preserving diverse image content. For example, computer-or AI-generated images may appear to provide suitable unreal counterparts to real images. However, making a well-controlled comparison, particularly at the neural level, is extremely difficult because numerous image features may vary simultaneously in ways that are difficult to characterize or control. Even when comparing natural images with corresponding realistic paintings, it is difficult to fully control for differences in factors such as image statistics and spatial layout. We therefore focused on Neural Style Transfer (NST), a technique developed in recent computer vision research^23^. By separately treating image content and surface style statistics in CNN feature space, NST enables systematic modulation of surface style while largely preserving the content of natural images. Indeed, NST has also been used in cognitive psychological studies investigating style perception and the contribution of style features to perception^24,25^. In the present study, we therefore used NST to generate unreal images with reduced image realism, and further constructed stimulus sequences in which perceived realness varied gradually by continuously morphing at the pixel level between each real image and its corresponding unreal image (Fig. 1b).

In the present study, to elucidate the neural basis of perceived realness, we conducted fMRI and MEG experiments in which participants viewed a wide variety of natural images (real images) and corresponding unreal images with modulated image realism. We found that the perceived realness of visual stimuli was prominently predicted by activity in higher dorsal visual areas, even after removing components explainable by image-computable features and decision-related activity. Representational similarity analysis further demonstrated that neural representations were transformed along the dorsal visual stream, from image-feature-based representations to perceived-realness-based representations. Finally, time-resolved source-level MEG decoding showed that the prediction peak in higher dorsal visual areas emerged later than those in early visual areas and lower-order dorsal visual areas.

Our results indicate that the human visual system first encodes image-computable features related to image realism, which are subsequently transformed into a representation of perceived realness along the dorsal visual stream. These findings further suggest that perceived realness is encoded at higher stages of the dorsal visual stream as an intermediate representation that bridges visual information and the potential for action upon and interaction with visual stimuli.

## Results

In the present study, we measured brain activity using two complementary stimulus sets generated with Neural Style Transfer (NST)^23^. In fMRI experiments 1–1 and 1–2, we used an image set in which image realism was parametrically varied across content-matched images, with the aim of minimizing unintended variation in other image features (Fig. 1b). This allowed us to investigate brain regions whose activity reflected graded changes in perceived realness. Specifically, we examined neural representations of perceived realness beyond components explainable by image features and decision-related activity. In fMRI experiment 2, we used a larger and more diverse image set containing real images and NST-synthesized images (Fig. 1c), with the aim of robustly identifying brain regions representing image features. In the MEG experiment, we also used the larger stimulus set (Fig. 1c) to reliably estimate source-localized MEG activity.

To investigate changes in perceived-realness-related information processing across visual areas while avoiding the risk of circular analysis^26^, we restricted our analyses a priori to retinotopically and functionally defined visual cortex (V1, V2, V3, hV4, VO, PPA, VTC, V3A, V3B, LO, hMT+, V7, IPS1, IPS2, and IPS3–IPS5; Fig. 2). To accurately account for individual differences in retinotopic organization, we conducted retinotopic and functional localizer scans for all observers who participated in the fMRI and MEG experiments and individually identified all visual areas. Hereafter, when referring to broader groups of visual areas, we use the following definitions: early visual areas, V1, V2, and V3; ventral visual areas, hV4, VO, PPA, and VTC; lateral visual areas, LO and hMT+; early dorsal visual areas, V3A and V3B; and higher dorsal visual areas, V7, IPS1, IPS2, and IPS3–IPS5.

**Figure 2.**
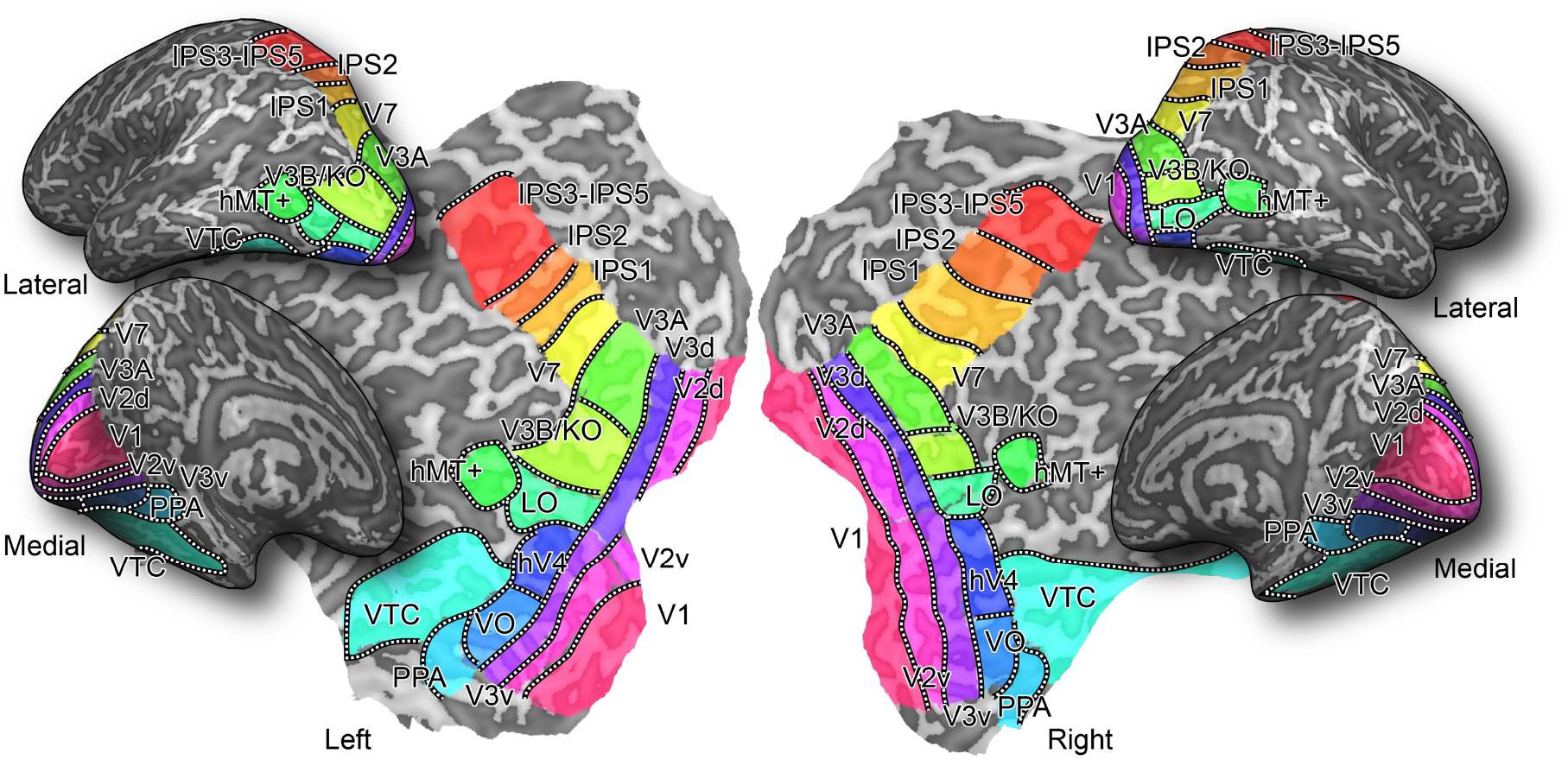
Regions of interest. We investigated neural responses in predefined regions of interest (ROIs) in the human visual cortex. ROIs were identified individually for each participant using separate retinotopic and functional localizer scans. The identified ROIs comprised V1, V2, and V3 as early visual areas; hV4, VO, PPA, and VTC as ventral visual areas; LO and hMT+ as lateral visual areas; V3A and V3B as early dorsal visual areas; and V7, IPS1, IPS2, and IPS3–IPS5 as higher dorsal visual areas. Adjacent areas that could not be reliably separated because of low signal-to-noise ratio were combined into a single ROI: VO comprised VO1 and VO2, LO comprised LO1 and LO2, and IPS3–IPS5 comprised IPS3, IPS4, and IPS5.

### Graded modulation of perceived realness

In fMRI experiment 1–1, on each trial, participants viewed images with graded levels of image realism (Fig. 1b) and rated their perceived realness (i.e., how much each visual stimulus appeared to capture the real world) on a four-point scale: “completely real,” “somewhat real,” “somewhat unreal,” and “completely unreal” (Fig. 1d). As shown in Fig. 1e, the mean ratings across 25 participants changed systematically with morph level (linear mixed-effects model: β = 0.297, 95 % CI = 0.293–0.300, *P* < 1.0 × 10^-100^). These results confirmed that the manipulation of image realism using NST and pixel-level linear interpolation induced graded changes in perceived realness.

### Perceived realness was best predicted in higher dorsal visual areas

We next decoded the perceived-realness response on each trial from the multivoxel BOLD pattern in each ROI. Perceived realness was predicted throughout the retinotopically defined visual cortex, but was particularly strongly represented in IPS1–IPS5 (Fig. 3b; all FDR-adjusted *P* < 0.05). Balanced accuracy in higher dorsal visual areas was significantly higher than that in early visual areas (FDR-adjusted *P* = 6.1 × 10^-8^, *t*(24) = 7.71, mean difference in balanced accuracy (ΔBA) = 6.09 %, 95 % CI [4.46, 7.72] %), ventral visual areas (FDR-adjusted *P* = 9.8 × 10^-9^, *t*(24) = 9.12, ΔBA = 6.72 %, 95 % CI [5.20, 8.24] %), early dorsal visual areas (FDR-adjusted *P* = 3.4 × 10^-8^, *t*(24) = 8.10, ΔBA = 5.15 %, 95 % CI [3.84, 6.46] %), and lateral visual areas (FDR-adjusted *P* = 9.8 × 10^-9^, *t*(24) = 8.86, ΔBA = 6.69 %, 95 % CI [5.13, 8.25] %).

**Figure 3.**
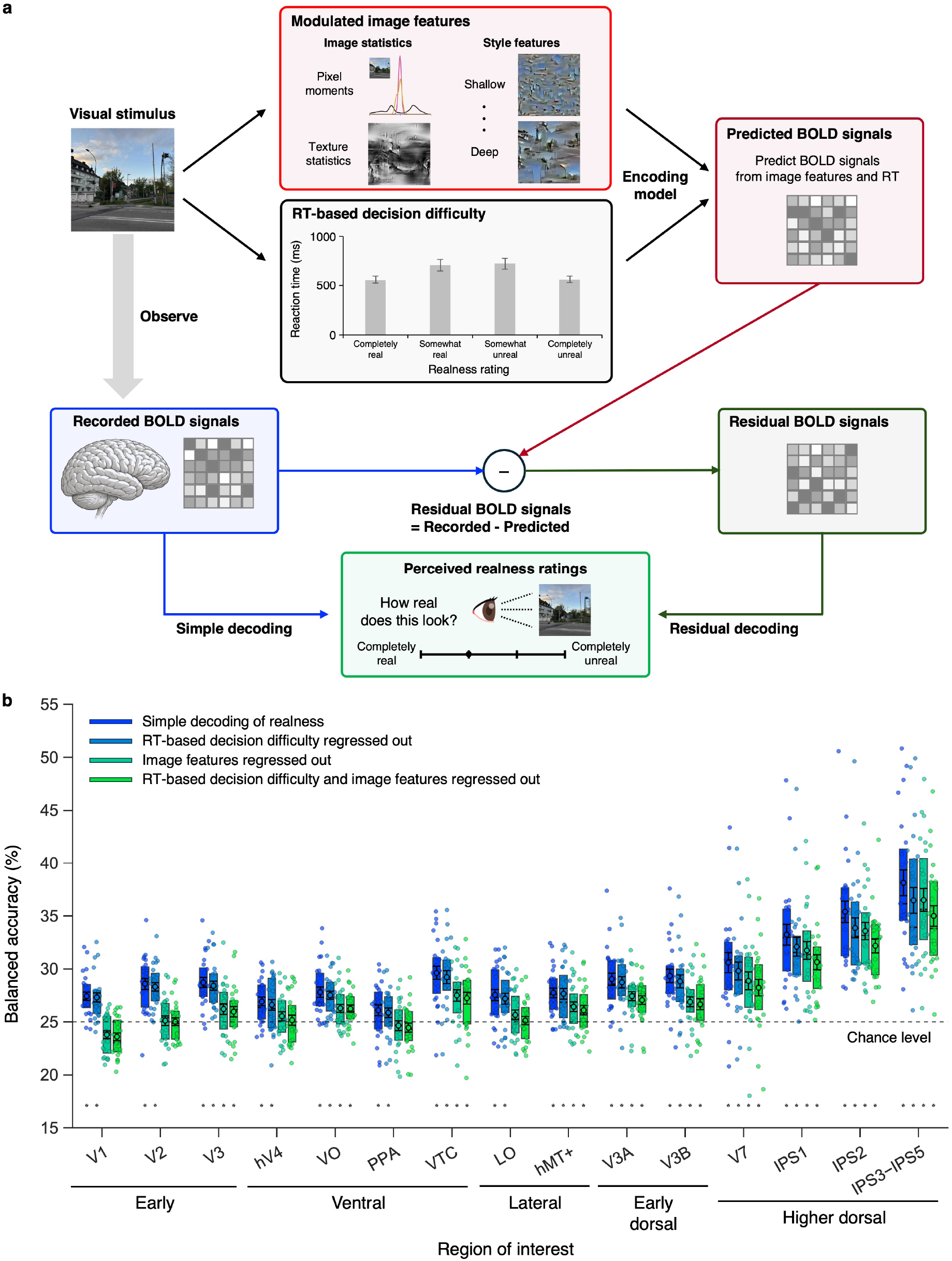
fMRI residual decoding of perceived realness. **(a)** Overview of the residual decoding analysis. We computed image features that varied between real and unreal images and measured reaction time (RT) on each trial as a proxy for decision difficulty. We then constructed encoding models to predict BOLD signals from these factors. Residual BOLD signals were obtained by subtracting the BOLD signals predicted by the encoding models from the observed BOLD signals, thereby removing variance explainable by these factors. **(b)** Decoding results for each condition. For each ROI, the four sets of boxes from left to right show balanced accuracy for simple decoding, decoding after removing RT-based decision difficulty, decoding after removing image-feature-related components, and decoding after removing both components. Dots indicate balanced accuracy for individual participants. Boxes extend from the first to the third quartile, with horizontal lines indicating the median. Black-outlined circles with error bars indicate the mean ±1 s.e.m. across participants. Asterisks below the boxes indicate statistically significant decoding relative to chance level (25 %; one-sided one-sample *t*-tests, FDR-adjusted *P* < 0.05, corrected across all ROIs and conditions).

These results cannot be explained solely by contralateral motor activity associated with right-hand button presses. Although all participants responded with their right hand in fMRI experiment 1–1, prediction accuracy was not consistently higher in the left hemisphere, which is contralateral to right-hand movements. Instead, balanced accuracy was relatively higher in the right hemisphere for IPS1 and in the left hemisphere for IPS2 and IPS3–IPS5. Moreover, significant prediction was consistently observed in right-hemisphere IPS areas (Supplementary Fig. 1a; all FDR-adjusted *P* < 0.05).

### Perceived realness was best predicted from activity in higher dorsal visual areas even after controlling for image features and nonspecific decision-related signals

Although the prominent decoding performance in higher dorsal visual areas might not necessarily reflect perceived realness itself, our additional analyses showed that it could not be explained solely by the image features that varied with image realism or by nonspecific processes involved in making responses.

If prediction in IPS areas were driven by image features that covaried with perceived realness or by decision-related components unrelated to perceived realness itself, no information about perceived realness should remain in the residual BOLD signals after removing the variance explainable by these components. In addition, if prediction in IPS areas were entirely driven by decision-related components, prediction accuracy should remain unaffected even after removing the variance explainable by image features. We therefore performed residual decoding^27^ (Fig. 3a) under the following three conditions and compared balanced accuracy:

1. Decoding after removing image-feature-related components
2. Decoding after removing reaction-time-(RT-) based decision difficulty
3. Decoding after removing both image-feature-related components and RT-based decision difficulty

Here, we considered two classes of image-computable features: image statistics based on the Portilla–Simoncelli model^28^, which also varied as a consequence of NST, and style features that were explicitly manipulated by NST.

Fig. 3b shows the results of the residual decoding analyses. Both RT-based decision difficulty and image features significantly reduced decoding accuracy in higher dorsal visual areas, indicating that both factors contributed to the decoding results in IPS areas. Nevertheless, balanced accuracy in higher dorsal visual areas remained significantly higher than that in the other visual areas under all residual conditions (all FDR-adjusted *P* < 0.05), demonstrating that perceived-realness-related information in higher dorsal visual areas remained even after controlling for image features and RT-related factors.

In detail, removing RT-based decision difficulty significantly reduced prediction accuracy in V3B and higher dorsal visual areas (FDR-adjusted *P* < 0.05). However, significant balanced accuracy was maintained in all visual areas even after removing the component related to RT-based decision difficulty (all FDR-adjusted *P* < 0.05). Thus, although perceived-realness decoding in higher dorsal visual areas may be partially explained by purely decision-related activity, decoding performance remained highest even after removing this component.

In contrast, removing the image-feature-related component significantly reduced balanced accuracy in all ROIs (all FDR-adjusted *P* < 0.05 across all ROIs). However, this reduction was smaller in higher dorsal visual areas, where significant residual decoding remained and was higher than in the other visual areas (FDR-adjusted *P* < 0.05). Furthermore, even after removing both image features and RT-related signals, perceived-realness responses were significantly predicted in higher dorsal visual areas, including IPS1–IPS5, with particularly strong decoding performance (FDR-adjusted *P* = 9.1 × 10^-8^, *t*(24) = 7.90, ΔBA = 5.63 %, 95 % CI [4.16, 7.10] % for IPS1; FDR-adjusted *P* = 1.2 × 10^-9^, *t*(24) = 10.9, ΔBA = 7.17 %, 95 % CI [5.81, 8.53] % for IPS2; FDR-adjusted *P* = 1.7 × 10^-9^, *t*(24) = 10.3, ΔBA = 10.0 %, 95 % CI [7.99, 12.0] % for IPS3−IPS5). As shown in Supplementary Fig. 2, the encoding models provided statistically significant fits in most ROIs, and the decoding accuracy for the corresponding features was indeed significantly reduced after residualization.

Together, these results demonstrate that perceived-realness decoding in higher dorsal visual areas cannot be explained solely by image features that vary with image realism or by purely decision-related information associated with the rating response.

### Decoding in higher dorsal visual areas was replicated after changing the rating task

As shown in Supplementary Fig. 1b, we further found that perceived-realness decoding in higher dorsal visual areas could not be explained by the selection among four response alternatives or by graded magnitude estimation. In fMRI experiment 1–2, while using the same stimulus set, participants judged each image as either a “photograph” or a “painting” and did not perform the four-level realness rating. The perceived realness obtained in fMRI experiment 1–1 was decoded from the BOLD signals measured under this task (all FDR-adjusted *P* < 0.05).

These results indicate that the information in higher dorsal visual areas cannot be explained solely by the four-button response mapping specific to fMRI experiment 1–1 or by the process of explicitly estimating the magnitude of perceived realness on a four-level scale.

### Image-computable features were broadly represented across early visual, ventral visual, and early dorsal visual areas

Fig. 4 shows which ROIs encoded image features that varied with image realism. As hypothesized, these image features were prominently encoded in early visual areas and ventral visual areas. Such image features were also encoded in early dorsal visual areas, whereas image-feature decoding accuracy was relatively low in higher dorsal visual areas, where perceived-realness decoding performance was highest.

**Figure 4.**
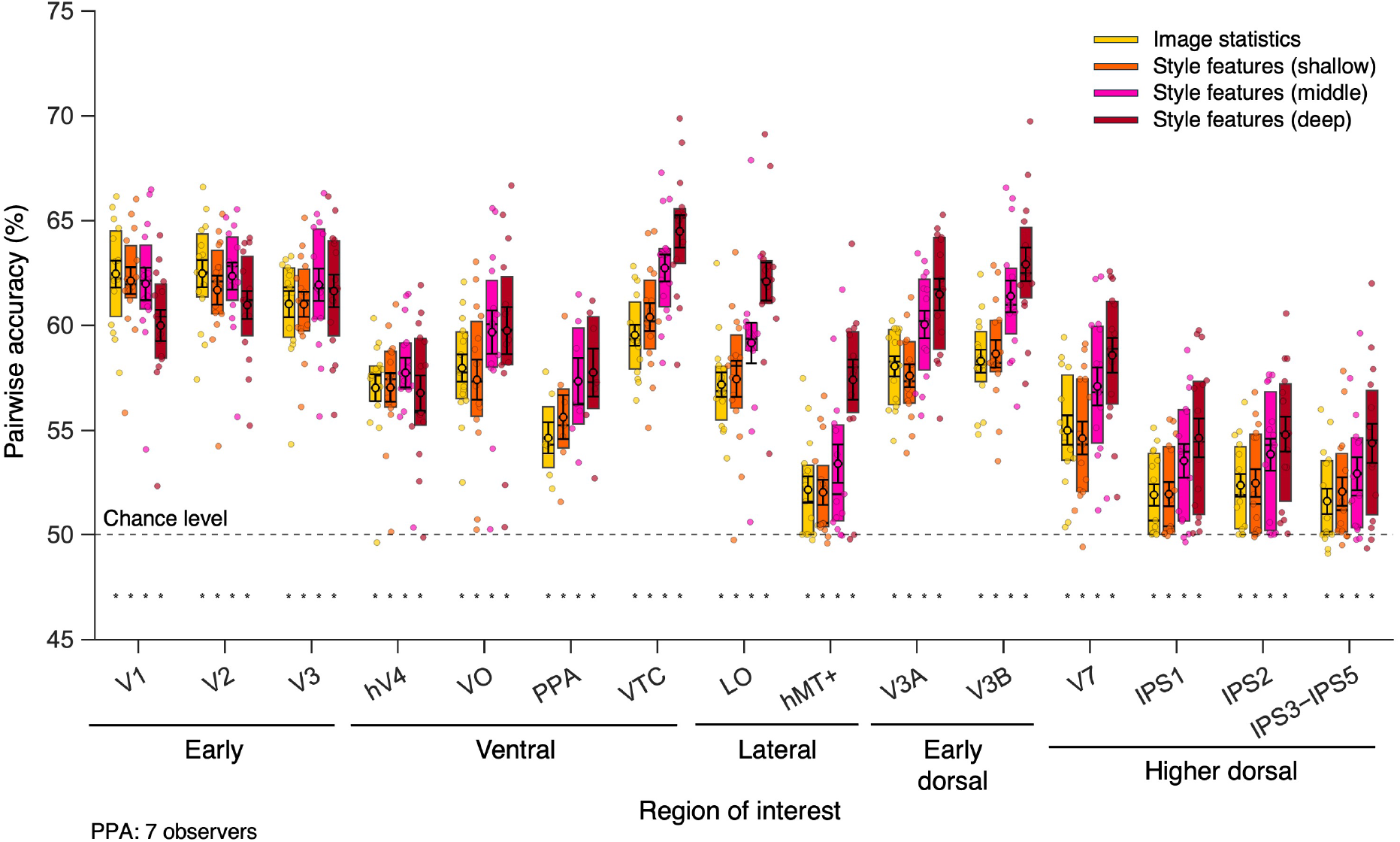
fMRI decoding of image features. Decoding results for each class of image features. For each ROI, the boxes from left to right show pairwise accuracy for image statistics and style features extracted from shallow, middle, and deep layers. Dots indicate pairwise accuracy for individual participants. Boxes extend from the first to the third quartile, with horizontal lines indicating the median. Black-outlined circles with error bars indicate the mean ±1 s.e.m. across participants. Asterisks below the boxes indicate statistically significant decoding relative to the theoretical chance level for pairwise accuracy (50 %; FDR-adjusted *P* < 0.05).

In fMRI experiment 2, we measured BOLD responses while participants viewed a total of 6,900 real images and 6,900 corresponding NST-synthesized unreal images. Because the stimuli used in fMRI experiments 1–1 and 1–2 were designed to minimize changes in image features, a stimulus set with greater variance in image features was required to reliably assess which visual areas represented each type of image feature. All image features were represented at statistically significant levels across all ROIs (df = 14 for all ROIs except PPA, for which df = 6; all FDR-adjusted *P* < 0.05), although prediction accuracy varied across ROIs. Image statistics computed based on Portilla–Simoncelli statistics^28^ were predicted with particularly high pairwise accuracy from early visual areas. In contrast, mid-to deep-layer style features from VGG19 were predicted significantly and with high accuracy from higher ventral visual areas, as well as from V3A, V3B, and LO. In higher dorsal visual areas, all classes of image features were predicted at statistically significant levels, but prediction accuracy was relatively low.

These results suggest that image-computable information, ranging from low-level image statistics to higher-level style features, that distinguishes unreal images from real images is prominently represented across early visual, ventral visual, and early dorsal visual areas, whereas it is represented only weakly in higher dorsal visual areas.

### Neural representations shifted from image-feature-based to perceived-realness-based

The decoding results suggested a progression in which image features that characterize image realism, ranging from low-to high-level features, are encoded across early visual and early dorsal visual areas and subsequently transformed into a representation of perceived realness in higher dorsal visual areas. We next used representational similarity analysis (RSA; Fig. 5a) to examine this transformation and found that neural representations shifted from image-feature-based to perceived-realness-based from early visual areas to higher dorsal visual areas.

**Figure 5.**
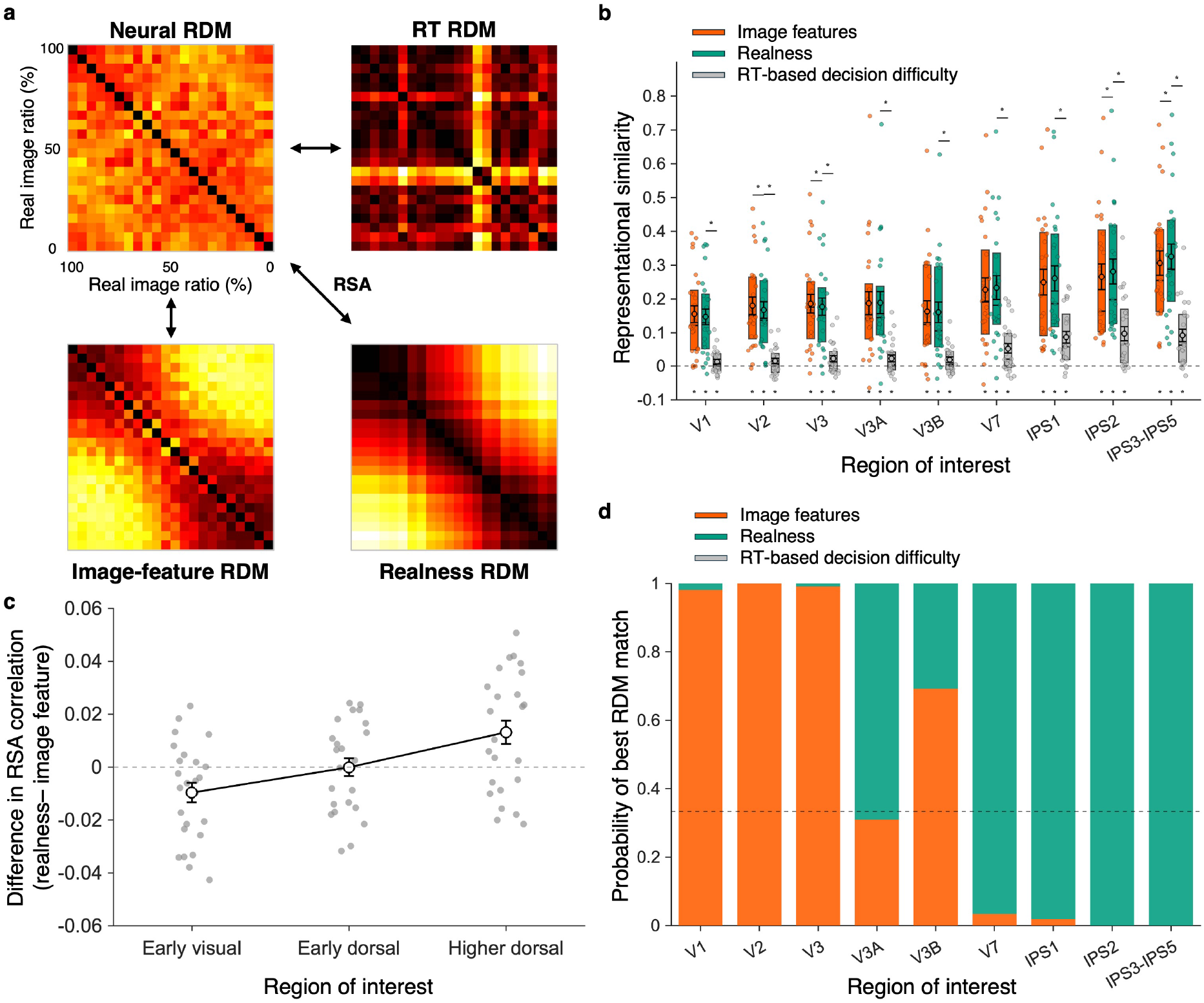
Results of representational similarity analysis. **(a)** Overview of the representational similarity analysis (RSA). We computed representational dissimilarity matrices (RDMs) for BOLD signals in each ROI (neural RDMs), image features (image statistics and style features), perceived realness, and RT-based decision difficulty. We then calculated Spearman correlations between the neural RDM in each ROI and each model RDM. **(b)** Correlations between neural and model RDMs. For each ROI, the boxes from left to right show correlations with the image-feature RDM, perceived-realness RDM, and RT RDM. Dots indicate individual participants. Boxes extend from the first to the third quartile, with horizontal lines indicating the median. Black-outlined circles with error bars indicate the mean ±1 s.e.m. across participants. Asterisks below the boxes indicate positive correlations that were significantly greater than expected under the permutation null distribution (one-sided permutation tests, FDR-adjusted *P* < 0.05). Asterisks above the boxes indicate statistically significant differences between model correlations. **(c)** Difference between the correlations with the perceived-realness RDM and the image-feature RDM (perceived realness − image features) for each ROI group. Dots indicate values for individual participants. **(d)** Probability that each model showed the highest similarity to the neural RDM. Bars indicate the probability that each model was the best-fitting model in each ROI, estimated using bootstrap resampling of participants.

As shown in Fig. 5b, image-feature RDMs showed high correlations with neural RDMs in early visual areas. In early visual areas (V2 and V3), the correlations between the neural RDMs and image-feature RDMs were significantly higher than those between the neural RDMs and the perceived-realness RDM (FDR-adjusted *P* = 5.0 × 10^-3^, |*t*(24)| = 3.33, Δρ = −0.012, 95 % CI [−0.0201, −0.00473] in V2; FDR-adjusted *P* = 4.6 × 10^-2^, |*t*(24)| = 2.26, Δρ = −0.008, 95 % CI [−0.0161, −7.19 × 10^-4^] in V3). In contrast, correlations with the perceived-realness RDM increased along the dorsal visual hierarchy, and the perceived-realness model showed the highest correlations in higher dorsal visual areas (Fig. 5b). In higher dorsal visual areas (IPS2 and IPS3–IPS5), the correlations between the neural RDMs and the perceived-realness RDM were significantly higher than those between the neural RDMs and image-feature RDMs (FDR-adjusted *P* = 7.9 × 10^-3^, *t*(24) = 3.07, Δρ = 0.016, 95 % CI [0.00510, 0.0260] in IPS2; FDR-adjusted *P* = 7.9 × 10^-3^, *t*(24) = 3.08, Δρ = 0.019, 95 % CI [0.00631, 0.0318] in IPS3−IPS5).

Moreover, as shown in Fig. 5c, this shift from image-feature-based to perceived-realness-based representations was statistically significant. The difference between the RSA correlations for the perceived-realness and image-feature models increased significantly along the dorsal visual hierarchy (one-sided *P* = 3.6 × 10^-5^, *t*(24) = 4.78, mean slope = 0.0114, 95 % CI [0.0065, 0.0163]).

This trend was also consistent with the results of a model comparison identifying the model that best explained the neural RDM in each ROI (Fig. 5d). To assess how robust the best-fitting model for each ROI was to variation in the participant sample, we performed a bootstrap model comparison with participants as the resampling unit. In early visual areas, the image-feature model showed a high winner probability, whereas the winner probability of the perceived-realness model increased in early dorsal visual areas, and the perceived-realness model had the highest probability of being the winner in higher dorsal visual areas. This result indicates that, although image features and perceived realness were correlated within the stimulus space, the model that best explained the neural representations changed from an image-feature-based model in early visual areas to a perceived-realness-based model in higher dorsal visual areas.

Together, these analyses revealed that neural representations transformed from image-feature-based to perceived-realness-based along the dorsal visual stream.

### Realness-related information emerged later in higher dorsal visual areas

If neural representations shift from image-feature-based to perceived-realness-based along the dorsal visual stream, responses related to perceived realness in higher dorsal visual areas should emerge later than those in early visual areas and early dorsal visual areas. Consistent with this idea, the MEG decoding results shown in Fig. 6a indicated that perceived-realness-related information was encoded at earlier latencies in early visual areas and early dorsal visual areas, but at later latencies in higher dorsal visual areas.

**Figure 6.**
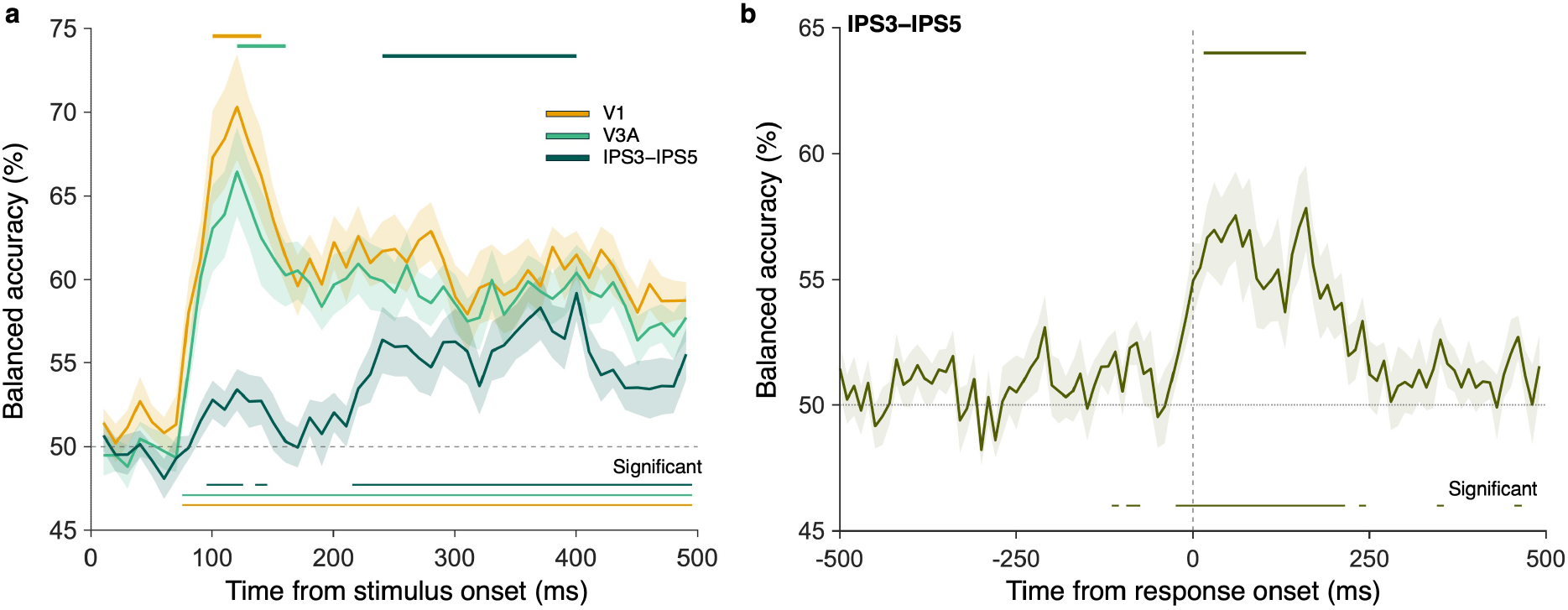
MEG decoding results. **(a)** Temporal profile of decoding accuracy for perceived realness from stimulus-locked, source-localized MEG signals. The yellow line indicates V1, the green line V3A, and the dark green line IPS3–IPS5. Bars below the plots indicate time points at which decoding accuracy was significantly above chance level (50 %; FDR-adjusted *P* < 0.05). Thick bars above the plots indicate peak-latency estimates obtained by participant bootstrap. **(b)** Temporal profile of response decoding accuracy from response-locked, source-localized MEG signals in IPS3–IPS5. Bars below the plot indicate time points at which decoding accuracy was significantly above chance level (50 %; FDR-adjusted *P* < 0.05). The thick bar above the plot indicates the peak-latency estimate obtained by participant bootstrap.

Decoding in the early visual cortex (V1) reached its maximum accuracy in the 111– 130 ms time window after stimulus onset, corresponding to a peak latency of 120.5 ms (participant-bootstrap 95 % CI [100.5, 140.5] ms). In the early dorsal visual area V3A, decoding accuracy peaked in the 111–130 ms time window, with a peak latency of 120.5 ms (95 % CI [120.5, 160.5] ms). In contrast, in higher dorsal visual areas (IPS3–IPS5), decoding performance peaked later, in the 391–410 ms time window, corresponding to a peak latency of 400.5 ms (95 % CI [240.5, 400.5] ms). Statistical significance of time-resolved decoding was assessed using one-sided one-sample *t*-tests against chance (FDR-adjusted *P* < 0.05 across all the plotted ROIs and all the time-windows). Peak latency was significantly later in IPS3–IPS5 than in V1 (FDR-adjusted *P* < 1.0 × 10^-4^, difference = 280 ms, 95 % CI [120, 300] ms) and V3A (FDR-adjusted *P* = 2.6 × 10^-3^, difference = 280 ms, 95 % CI [120, 280] ms), as assessed from the participant-bootstrap distributions of the peak-latency differences.

### Realness-related information in higher dorsal visual areas could not be explained solely by image-feature-related or response-related activity

The later processing of perceived-realness-related information in higher dorsal visual areas (IPS3–IPS5) could not be accounted for solely by image-feature processing or response-related activity. If it entirely reflected image-feature processing, the relevant image features should also be prominently decoded at similar latencies. If it reflected response-related activity, the peak of response-locked decoding should overlap with the peak of stimulus-locked decoding and should be temporally continuous with it. Our additional analyses supported neither possibility.

Supplementary Fig. 3 shows the results of predicting image features from stimulus-locked time-resolved source activity. Image statistics and shallow-, mid-, and deep-layer style features were first continuously predicted at statistically significant levels from 71–90 ms, 71−90 ms, 81−100 ms, 81−100 ms after stimulus onset in early visual areas (V1; FDR-adjusted *P* < 0.05 across all the time-windows and image feature classes) and from 81–100 ms, 71−90 ms, 71−90 ms, 71−90 ms in early dorsal visual areas (V3A; FDR-adjusted *P* < 0.05 across all the time-windows and image feature classes). In contrast, in higher dorsal visual areas (IPS3–IPS5), image features were not decoded at statistically significant levels across consecutive time windows (FDR-corrected across all time windows and image-feature classes in IPS3–IPS5). These results suggest that the prediction of perceived realness in higher dorsal visual areas did not simply reflect the encoding of image features.

Fig. 6b shows response-locked time-resolved decoding of real versus unreal images from source activity in higher dorsal visual areas (IPS3–IPS5). The group-average response-locked decoding curve reached its maximum at 160.5 ms after response onset, with a participant-bootstrap 95 % confidence interval of 15.5–160.5 ms after response onset. A one-sided participant-bootstrap test showed that the peak occurred significantly after response onset (*P* < 1.0 × 10^-4^). Importantly, behavioral responses occurred only after stimulus offset, whereas the stimulus-locked decoding peak in IPS3–IPS5 occurred nearly 100 ms before stimulus offset (Fig. 6a). Thus, the response-locked peak necessarily occurred later than the stimulus-locked peak, showing a temporal separation between the peaks identified in the stimulus-locked and response-locked analyses.

## Discussion

The present study revealed how the visual system transforms image features in visual stimuli into a representation of perceived realness by combining behavioural ratings, fMRI, and MEG. Image statistics and style features that characterize the differences between real and unreal images were represented across early visual areas, ventral visual areas, and early dorsal visual areas. In contrast, even after removing components explainable by these image-computable features and decision-related activity, perceived-realness-related information remained selectively in higher dorsal visual areas, particularly in IPS areas. Furthermore, the model that best explained the neural representation in each region shifted from an image-feature-based model to a perceived-realness-based model along the dorsal visual hierarchy. Time-resolved MEG analyses also showed that perceived-realness-related information emerged early in early visual areas and early dorsal visual areas, whereas it emerged later in higher dorsal visual areas. These results suggest that image-computable features that characterize image realism are processed hierarchically and transformed into a higher-level perceptual representation of perceived realness in higher dorsal visual areas.

The representation of image statistics in early visual areas is consistent with many previous studies showing the encoding of local luminance, orientation, spatial frequency, and subband statistics in early visual cortex^7,29–31^. Because low-level statistics such as edges and spatial-frequency power also varied systematically between real and unreal images, responses in early visual areas likely reflect basic visual cues that contribute to perceived realness. Style features, which also varied systematically between real and unreal images, contain hierarchical visual information ranging from local color and texture to shape- and object-related information^25^, and were therefore expected to be strongly represented in ventral visual areas that process such information^19,32–35^. Consistent with this prediction, mid-to deep-layer style features were well predicted from activity in higher ventral visual areas. At the same time, the finding that the style features were also encoded in early dorsal visual areas indicates that the processing of image information that characterizes image realism is not restricted to the ventral visual stream. Early dorsal visual areas have been reported to contribute not only to the processing of binocular disparity and three-dimensional structure^36–42^, but also to surface-related information such as perceived surface gloss^43,44^. In fact, anatomical connections between ventral and dorsal visual areas, the vertical occipital fasciculus connecting hV4 and V3A^45,46^, could provide a neural substrate for such distributed representations. Thus, the present study shows that style features, whose neural basis has not been well established in previous studies, are represented not only in ventral visual areas as expected, but also across early dorsal visual areas. These findings provide new insights into both the neural representation of visual style and the neural circuitry supporting distributed visual processing.

A central finding of the present study is that perceived-realness-related information remained in IPS areas even after removing components explainable by image-computable features and decision-related activity, and that neural representations shifted from image-feature-aligned to perceived-realness-aligned along the dorsal visual stream. These results suggest that perceived realness is not simply read out as the sum of image features that characterize image realism, but is instead formed as a higher-order representation that integrates these features. IPS areas, which carried neural representations of perceived realness, are broadly involved in processes that link visual information to space and action in the physical world, including the processing of three-dimensional structure^38,47^, spatial attention^48,49^, body-centered spatial representations^50,51^, and action-related processing involved in reaching and grasping^52,53^. In particular, studies comparing real three-dimensional objects with their two-dimensional images have shown that the neural processing of these stimulus formats can differ not only in the ventral visual stream but also in dorsal and parietal regions involved in visuomotor processing^54–56^. Given these functions, perceived-realness-related representations in IPS may reflect not only the degree to which a visual stimulus faithfully reproduces the appearance of the real world, but also whether the stimulus can be interpreted as part of a coherent physical world and the extent to which it can be accepted as a potential target for perception and action in the real world. In this sense, perceived realness may constitute an intermediate representation that emerges as the brain evaluates the potential for action upon and interaction with a visual stimulus, and may be encoded at a late stage of visual processing.

This interpretation of perceived realness highlights the need to distinguish the physical and statistical fidelity of an image from the perceived realness experienced by observers. In the context of computer graphics and generative AI, realness is often approached in terms of visual or physical fidelity, that is, how accurately optical behavior, local texture, shape, illumination, and image statistics reproduce those of the real world^2,57,58^. However, our finding that perceived-realness-related information emerged in higher dorsal visual areas in a manner distinct from the representation of image features suggests that physical fidelity alone may not fully account for perceived realness. First, even an image with extremely high physical fidelity does not necessarily appear real. For example, even in high-fidelity CG or AI-generated images, subtle inconsistencies among cues related to object structure, eye and facial movements, illumination, and physical contact can evoke a strong sense of incongruity or eeriness^4^. The uncanny valley in human-like stimuli^5,6^ provides a representative example: even when individual local features are reproduced with high fidelity, perceived realness can decrease if those features are not mutually coherent as a whole. Our findings provide a framework for understanding such phenomena not simply as a deficiency in particular image features, but as a failure of higher-order processing to integrate those features into a coherent representation of the real world. Second, a strong sense of realness can arise even when the physical fidelity of an image is low. Even in low-resolution VR environments with simple rendering, observers can experience an environment as highly real when changes in viewpoint, body movements, spatial layout, and responses of objects are mutually consistent^59–61^. Such experiences suggest that perceived realness is determined not only by the fidelity of an image, but also by the extent to which visual input can be incorporated into a coherent spatial and bodily representation of the world. Perceived-realness-related representations in higher dorsal visual areas may provide a neural basis for understanding this dissociation between visual fidelity and experienced realness. Thus, perceived realness in humans may need to be understood more broadly, not simply as how closely a stimulus resembles the real world, but as the extent to which the brain can interpret the stimulus as a coherent target of perception and interaction within the real world.

IPS is also involved in perceptual decision-making^62–67^, raising the possibility that the information observed here reflected decision-related activity associated with explicit ratings. However, perceived-realness-related information was replicated across fMRI experiments using different task conditions and remained after removing components explainable by RT-based decision difficulty. Thus, representations in higher dorsal visual areas cannot be explained, at least, solely by rating difficulty or simple decision-related demands. Instead, IPS may be involved in a stage at which distributed image features are read out into a representation of perceived realness that can be used for perceptual judgments and subsequent actions.

The perceived realness examined in the present study differs from reality monitoring, which involves judging whether a perceptual experience originates from external sensory input or is internally generated through processes such as imagery or memory. However, the two share the distinction between sensory representations themselves and higher-order processes that interpret those representations as real. Recent studies of reality monitoring have shown that reality judgments depend not only on the presence of sensory activity, but also on how higher-order regions read out the strength and reliability of that activity^68,69^. Similarly, in the present study, visual cues to image realism were distributed across early visual, ventral visual, and early dorsal visual areas, whereas perceived-realness-related representations emerged more clearly in higher dorsal visual areas. Together, these findings suggest that different forms of reality-related perception may rely on a common computational principle in which sensory evidence is transformed into a higher-order representation and interpreted as belonging to the real world.

Overall, the present study showed that image features related to image realism are broadly represented across the visual system, while being integrated and transformed into a higher-order perceptual representation of perceived realness in higher dorsal visual areas. These findings indicate that perceived realness is not determined solely by the physical fidelity of an image, but emerges from the process by which multiple sources of visual evidence are interpreted as a coherent representation of the real world.

## Materials & Methods

### Participants

#### fMRI experiment 1−1

Twenty-eight naïve human participants of either sex participated in the experiment. Their ages ranged from 21 to 28 years. The target sample size was determined a priori as 25–30 participants with reference to previous studies, with the aim of obtaining robust results across participants. Three participants who were unable to complete the experiment within the scheduled experimental period were excluded from the analysis.

#### fMRI experiment 1−2

Twenty naïve human participants of either sex participated in the experiment. Their ages ranged from 21 to 28 years. Participants were recruited, whenever possible, from those who had participated in fMRI experiment 1–1, with a target of at least 15 participants. Some participants completed fMRI experiment 1–2 before participating in fMRI experiment 1–1. Three participants were excluded from the analysis because they were unable to participate in fMRI experiment 1–1 due to scheduling constraints or because the experiment was interrupted before completion, leaving 17 participants for analysis.

#### fMRI experiment 2 and MEG experiment

Sixteen naïve human participants of either sex participated in fMRI experiment 2. Their ages ranged from 22 to 28 years. The initial target sample size of 13 participants was determined based on the number of trials required to present each of the 6,900 real images and the corresponding 6,900 NST-synthesized unreal images at least once. Three additional participants were recruited. Specifically, two participants were recruited as replacements for one participant who was unable to participate in the MEG experiment using the same visual stimuli because of scheduling constraints and one participant whose button-response rate was extremely low. The third additional participant was recruited to acquire BOLD data that had not been recorded because of the initial experimental protocol and thereby complete the dataset. The participant with the extremely low response rate was excluded from the fMRI experiment 2 analysis, leaving 15 participants for analysis. Because no corresponding data were missing in the MEG measurements, MEG data were acquired from 13 participants.

All participants had normal or corrected-to-normal vision. All ethical regulations relevant to research involving human participants were followed. All procedures were conducted in accordance with the relevant institutional guidelines and regulations and were approved by the Ethics and Safety Committees of the Center for Information and Neural Networks (CiNet), National Institute of Information and Communications Technology (NICT). Participants provided written and oral informed consent before the experiment. We attempted to recruit the same participants across experiments whenever possible; however, because the experiments were conducted over an extended period, a substantial number of participants were unable to participate in all experiments because of scheduling conflicts. In addition, all participants completed an fMRI localizer scan.

### Visual stimulus

#### Real images

The real images presented to participants during the experiments consisted of 6,900 natural scene images selected from the Microsoft Common Objects in Context (COCO) database^70^. The 6,900 real images were selected while excluding highly salient images, such as those containing many people or those in which a small number of large animals occupied a substantial portion of the image. The scatter plot in Fig. 1c shows the CLIP features of the real images after dimensionality reduction using t-SNE. As shown in the figure, deep features containing semantic information were reasonably dispersed across the visual stimuli. To avoid large differences in mean luminance across real images and the resulting differences in attention, the mean luminance of the real images was normalized to 0.5 times the maximum luminance. Because of camera gamma correction, the mean luminance of the images actually presented was 0.27 ± 0.03 times the maximum luminance, thereby reducing large variations in luminance across images. During stimulus presentation in the fMRI and MEG recordings, the luminance of the uniform background was 0.3 times the maximum luminance, which also prevented visual stimuli with substantially different brightness from strongly popping out.

#### Unreal images

We selected 1,600 paintings from the refined WikiArt dataset^71^ that were suitable for style transfer, excluding discrete images such as collections of geometric shapes. For each of the 6,900 real images, five paintings were randomly selected from the set of 1,600 paintings as style images, and neural style transfer (NST)^23^ was applied to generate five candidate unreal images. NST was implemented according to the PyTorch tutorial available as of February 2026 (https://docs.pytorch.org/tutorials/advanced/neural_style_tutorial.html), using VGG19 as the model, conv2_2 for the content loss, and conv1_1, conv1_2, conv2_1, conv2_2, and conv3_1 for the style loss, where convX_Y denotes the Y-th Conv2d layer in the X-th convolutional block. Although this choice of layers differed from that used in the original study^23^, optimizing the content loss at the relatively shallow conv2_2 layer was appropriate for the purpose of the present study, which was to modulate style while preserving image content. To avoid large differences in low-level image features and overall appearance between real images and their corresponding unreal images, each candidate image was further adjusted after NST so that the mean and standard deviation of each channel in L*a*b* color space matched those of the corresponding real image. Among the five candidates, the image with the smallest cosine distance in Portilla–Simoncelli image statistics from the corresponding real image was selected as the final unreal image.

#### Visual stimuli with graded levels of image realism

From the 6,900 real/unreal image pairs described above, 50 pairs of real and unreal images were selected. For each corresponding pair, intermediate images were generated by linearly combining the real and unreal images at ratios of 90:10, 80:20, …, and 10:90 at the pixel level (Fig. 1b). Together with the real image itself (100:0) and the unreal image itself (0:100), this yielded 11 stimuli with graded levels of image realism for each real image. The 50 image pairs used to generate these morphed images were selected in advance by 5 participants, including one of the authors, as pairs for which perceived realness appeared to change gradually across the morph continuum. Images that were not selected tended to show abrupt changes in perceived realness at particular morph levels and were therefore unsuitable for the purpose of the experiment.

### Image feature analysis

#### Image statistics computation

We computed image statistics equivalent to the Portilla– Simoncelli statistics^28^. Specifically, images were loaded after linearizing the image intensities, resized to 224 × 224 pixels, and converted from color to grayscale according to the NTSC standard. The resulting luminance images were decomposed into subband images at five spatial-frequency scales (112, 56, 28, 14, and 7 cycles/image) and eight orientations (0°, 22.5°, …, 157.5°). For each subband, we computed log(SD), skewness, log(kurtosis), and correlations across subbands, including cross-spatial-frequency and cross-orientation correlations. The resulting values were summarized by averaging across orientations^72^. To obtain statistics in CIELAB color space, the input images were converted using the MATLAB function rgb2lab, and the same procedure was applied separately to each channel. Following the previous study^19^, the image-statistics feature set consisted of subband statistics computed from the luminance images; the mean, log(SD), skewness, and log(kurtosis) of the L* channel in CIELAB space; and the mean and log(SD) of the a* and b* channels.

#### Style feature computation

We next computed the features optimized as the style loss in neural style transfer, namely the Gram matrices of intermediate activations in a deep convolutional neural network. Specifically, each image was fed into a pretrained VGG19 model, and channel-wise activation correlations were computed for layers conv1_1, conv1_2, conv2_1, conv2_2, conv3_1, conv4_1, and conv5_1, where convX_Y denotes the Y-th Conv2d layer in the X-th convolutional block. These layers included those used in the previous study^23^, together with conv1_2 and conv2_2, which were explicitly modulated during generation of the unreal images in the present study. Because concatenating all style features resulted in more than 600,000 feature dimensions, we reduced their dimensionality separately for each layer using truncated singular value decomposition (SVD), with the number of components selected according to a 95 % cumulative explained-variance threshold for each layer. For the large image dataset, which contained 13,800 samples, principal component analysis (PCA) was not performed; truncated SVD was used instead as a conceptually similar dimensionality-reduction procedure.

### Apparatus of fMRI/MEG experiments

#### fMRI experiments

Visual stimuli were generated on a PC (HP Z440 Workstation, HP Inc., California, USA) and presented on a gamma-corrected LCD display with a maximum luminance of 123 cd/m^2^ at a size of 768 × 768 pixels. At a viewing distance of 153 cm, this corresponded to a visual angle of 10° × 10°. All visual stimuli were treated as if they had been captured with a camera gamma of 0.5 and were loaded using a gamma value of 2.

#### MEG experiment

Visual stimuli were generated on a PC (Zotac QX3P5000, Zotac Technology Limited, Hong Kong, China) and presented via a PROPixx system (VPixx Technologies Inc., QC, Canada) on a screen with a maximum luminance of 191 cd/m^2^ at a size of 512 × 512 pixels. At a viewing distance of 50 cm, this corresponded to a visual angle of 10° × 10°. All visual stimuli were treated as if they had been captured with a camera gamma of 0.5 and were loaded using a gamma value of 2. Because the baseline luminance was too high, a 10 % neutral-density (ND) filter was used to reduce the luminance.

### Experimental procedures

#### fMRI experiment 1−1

Each experiment consisted of 11 runs, with 50 images presented in each run. At the beginning of each run, a uniform background was presented for 16 s, after which the first image was presented for 2 s. This was followed by a uniform background for either 4 or 6 s. Thereafter, images were presented sequentially for 2 s each, with an interstimulus interval (ISI) of either 4 or 6 s. After the final image disappeared, only the uniform background was presented for 16 s. This procedure resulted in a total of 550 trials. Across these 550 trials, each of the 550 images was presented once in a random order. To directly relate participants’ subjective evaluations of each visual stimulus to the corresponding neural activity recorded during scanning, we asked participants to rate its “realness” (“How real does this look?”) on a four-level scale after the image disappeared, using a button press with their right hand. Specifically, with a photograph defined as the most real, participants responded using one of four categories: “completely real,” “somewhat real,” “somewhat unreal,” and “completely unreal.” The mean responses across participants are shown in Fig. 1e.

#### fMRI experiment 1−2

The experimental procedure was identical to that of fMRI experiment 1–1 except for the response criterion. In fMRI experiment 1–2, after each stimulus disappeared, participants performed a categorical judgment of whether the stimulus was a “photograph” or a “painting.” Participants responded by pressing one of two buttons after each trial, thereby maintaining attention to the stimuli.

#### fMRI experiment 2

From the 6,900 real images, 400 images were selected as a common image set, and the remaining 6,500 images were divided into 13 sets of 500 images. Across scanning sessions conducted over 4 days, each participant viewed the 400 common images, 500 unique images, and their corresponding unreal images, yielding a total of 1,800 images, each presented once while fMRI data were acquired. Each scanning session consisted of 9 runs, with 50 images presented in each run. At the beginning of each run, a uniform background was presented for 16 s, after which the first image was presented for 2 s. This was followed by a uniform background for either 4 or 6 s. Thereafter, images were presented sequentially for 2 s each, with an ISI of either 4 or 6 s. Repeating this scanning procedure four times resulted in a total of 36 runs and 1,800 trials. Across these 1,800 trials, each of the 1,800 images was presented once in a random order. For some participant scans, the waiting period after disappearance of the final image in each run was too short to obtain a BOLD signal corresponding to that final image. Additional measurements were therefore conducted for some participants to compensate for these missing data, such that at least one BOLD response was obtained for each of the 13,800 images. In addition, sub-16 was newly recruited for these additional measurements. After each image disappeared, participants indicated by button press whether the image was real or unreal. Because performance on this task was nearly 100 % correct, we considered it suitable as an attention-control task while minimizing differences in attentional demands across stimuli.

A 10-s dummy scan was performed before each scan, and the resulting data were discarded. At the end of each day’s scanning session, we also acquired a top-up scan for image distortion correction.

#### MEG experiment

Participants who took part in fMRI experiment 2 viewed the same image set in the MEG experiment. This large set of real and unreal images allowed us to present many distinct images while collecting a large number of trials within the limited recording time available for MEG. The MEG experiment consisted of 16 runs conducted over two days (8 runs per day), with 225 images presented in each run. At the beginning of each run, a uniform background was presented for 1 s, followed by the first image for 0.5 s. Subsequent images were presented sequentially for 0.5 s each, with an interstimulus interval (ISI) of either 1 or 1.2 s, during which a uniform background was displayed. Each image was presented once per day, allowing brain responses to each image to be measured twice across the two days. After each image disappeared, participants indicated by button press whether the image was real or unreal. Because performance on this task was nearly 100 % correct, we considered it suitable as an attention-control task while minimizing differences in attentional demands across stimuli.

#### fMRI localizer scan

ROIs were delineated individually for each participant using standard retinotopic mapping procedures with a publicly available stimulus package (https://github.com/hiroshiban/Retinotopy). Retinotopic maps were obtained using rotating wedge stimuli presented in clockwise and counterclockwise directions, expanding and contracting concentric rings, and moving-bar stimuli. These maps were used to delineate the boundaries of V1, V2, V3, V3A, V3B, LO (LO1 and LO2), V7, IPS1, IPS2, IPS3–IPS5, hV4, VO (VO1 and VO2). Functional localizer scans were additionally conducted to identify hMT+ using moving versus stationary dot stimuli, V3B/KO using a kinetic-object localizer, and the parahippocampal place area (PPA) using the contrast between buildings and the other stimulus categories. The localization of V3B/KO was guided by previously reported anatomical and functional characteristics of the kinetic occipital region^40,73^. Results from these functional localizers were used as complementary information to support ROI delineation based primarily on the retinotopic maps. Adjacent areas whose boundaries could not be clearly distinguished based on the retinotopic maps were not forcibly separated but were instead combined into a single ROI for subsequent analyses, taking their functional similarity into account. Specifically, VO consisted of VO1 and VO2, LO consisted of LO1 and LO2, and IPS3–IPS5 consisted of IPS3, IPS4, and IPS5. PPA was defined based on the functional localizer, whereas VTC was defined with reference to the DKT atlas, following a procedure similar, but not identical, to that used in previous studies^74–76^. Both regions were identified so as not to overlap with any of the other ROIs.

### fMRI acquisition

#### T1-weighted images

A high-resolution T1-weighted anatomical image was acquired using a three-dimensional magnetization-prepared rapid gradient-echo (MPRAGE) sequence. The acquisition parameters were as follows: TR = 1,900 ms, TE = 2.26 ms, inversion time (TI) = 900 ms, flip angle = 9°, voxel size = 1 × 1 × 1 mm^3^, field of view = 256 × 256 mm^2^, and acquisition matrix = 256 × 256. Parallel imaging was performed using GRAPPA with an acceleration factor of 2. The anatomical images were acquired in the sagittal orientation.

#### fMRI experiment 1-1, 1-2, and 2

Functional MRI data were acquired using a 3-T Siemens PrismaFit MRI scanner (Siemens Healthineers, Erlangen, Germany) equipped with a 64-channel head/neck coil. Functional images were acquired using a T2*-weighted multiband gradient-echo echo-planar imaging (EPI) sequence provided by the Center for Magnetic Resonance Research (CMRR), University of Minnesota, with the following parameters: repetition time (TR) = 2,000 ms, echo time (TE) = 30 ms, flip angle = 75°, voxel size = 2 × 2 × 2 mm^3^, field of view = 192 × 192 mm^2^, acquisition matrix = 96 × 96, 62 slices, and multiband acceleration factor = 2. In-plane parallel imaging with an acceleration factor of 2 (GRAPPA) was also used. The phase-encoding direction was anterior-to-posterior. The effective echo spacing was 0.345 ms, and the total readout time was 32.8 ms.

#### Retinotopy mapping and functional localizer scans

For retinotopic mapping and functional localizer scans, functional images were acquired using a T2*-weighted multiband gradient-echo EPI sequence provided by the Center for Magnetic Resonance Research (CMRR), University of Minnesota. The acquisition parameters were as follows: repetition time (TR) = 1,000 ms, echo time (TE) = 30 ms, flip angle = 60°, voxel size = 2 × 2 × 2 mm^3^, field of view = 192 × 192 mm^2^, acquisition matrix = 96 × 96, 45 slices, and multiband acceleration factor = 3. In-plane parallel imaging was performed using GRAPPA with an acceleration factor of 2. The phase-encoding direction was anterior-to-posterior. The effective echo spacing was 0.345 ms, and the total readout time was 32.8 ms.

### fMRI preprocessing

#### Preprocessing

Results included in this manuscript come from preprocessing performed using fMRIPrep 23.2.3^77^ (RRID:SCR_016216), which is based on Nipype 1.8.6^78^ (RRID:SCR_002502). The preprocessed fMRI time series were additionally subjected to high-pass temporal filtering in BrainVoyager before GLM estimation and subsequent multivoxel analyses. Further details of the preprocessing procedures are provided in the Supplementary Methods.

#### General linear models

Voxels used for the subsequent prediction analyses were selected using a general linear model (GLM) implemented in BrainVoyager (Brain Innovation, Maastricht, The Netherlands). Before GLM estimation, the preprocessed fMRI time series were converted to percent signal change using BrainVoyager. For each participant, all functional runs were concatenated and entered into a voxel-wise GLM. Head motion parameters (3 translations and 3 rotations) and run-specific constant terms were additionally included as nuisance regressors. Regression coefficients (beta values) were estimated using ordinary least squares. To identify voxels responsive to visual stimulation, a contrast comparing all stimulus presentations with the fixation baseline was computed without distinguishing among the experimental conditions. Thus, voxel selection was based on the overall response to visual stimulation rather than on condition-specific response differences. Within each individually defined region of interest (ROI), voxels were ranked according to the resulting t-values, and the 500 voxels with the highest t-values were selected as features for the subsequent prediction analyses. For the subsequent prediction analyses, the preprocessed fMRI time series were z-score standardized separately for each voxel within each run. To obtain the BOLD response associated with each stimulus presentation, the standardized signal was averaged separately for each selected voxel over a time window from 4 to 10 s after stimulus onset. The resulting voxel-wise response pattern for each stimulus presentation was used in the subsequent analyses.

#### ROI definition

All participants who took part in the fMRI experiments also underwent standard retinotopic mapping and functional localizer scans, either in the same session or in a separate session. Retinotopic mapping was performed using the built-in procedures in BrainVoyager, with the BOLD time series expressed as percent signal change. Individual regions of interest (ROIs) were defined for each participant on the basis of these measurements. V1, V2, V3, hV4, V3A, V3B, V7, IPS1, and IPS2 were delineated individually on the basis of retinotopic maps following previous studies. Because the boundaries among IPS3, IPS4, and IPS5 were not clearly identifiable in some participants, these areas were combined and treated as a single ROI (IPS3–IPS5). The definition of V3B was additionally guided by responses obtained from the kinetic-object (KO) functional localizer. Because clear boundaries between VO1 and VO2 could not be identified consistently across participants, these regions were combined into a single VO ROI. VO was defined as the region medial to the lower-visual-field representation and the central visual-field representation used to delineate hV4. The region lateral to this boundary did not correspond well to the visual-field-map organization described in previous retinotopic studies and was therefore defined as ventral temporal cortex (VTC), following the previous studies^74–76^. The VTC ROI included part of the fusiform gyrus as defined by the Desikan–Killiany–Tourville (DKT) atlas. The human motion-sensitive complex (hMT+) was identified using an MT functional localizer. Because the boundary between LO1 and LO2 was also ambiguous in some participants, these regions were combined into a single lateral occipital (LO) ROI. LO was defined as the region ventral to V3B, posterior to hMT+, and dorsal to the foveal representation. Finally, the parahippocampal place area (PPA) was identified on the basis of responses obtained from the functional localizer, with voxels overlapping the VO ROI excluded.

### fMRI data analysis

#### Behavioural analysis

To assess the relationship between morph level and perceived-realness ratings, we fitted a linear mixed-effects model using the fitlme function in MATLAB, with morph level as a fixed effect and random intercepts for participants and original images. Trials with missing ratings were excluded from the analysis.

#### Perceived realness decoding

To test whether perceived realness could be decoded from fMRI activity recorded in fMRI experiment 1−1, we trained a four-class linear support vector machine (SVM) separately for each participant and ROI to predict the participant’s perceived-realness rating (0–3) from the voxel-wise BOLD response pattern. Multiclass classification was implemented in MATLAB using an error-correcting output codes (ECOC) model (fitcecoc) comprising one-versus-one linear SVM classifiers. Predictors were standardized during SVM training. Samples for which a valid perceived-realness response or BOLD response pattern was unavailable were excluded from the analysis. On average, 1.2 % of trials were excluded per participant (range: 0–14.0 %).

Prediction performance was evaluated using 10-fold cross-validation. To prevent classification from relying on image-specific information, the cross-validation folds were defined according to source-image identity. Specifically, all morph images derived from the same original image were assigned to the same fold, such that stimuli originating from a given source image never appeared in both the training and test sets. Source-image identities were partitioned into 10 non-overlapping folds; the classifier was trained on nine folds and tested on the remaining fold, and this procedure was repeated until each fold had served once as the test set.

Predicted labels from all cross-validation folds were pooled before decoding accuracy was calculated for each participant and ROI. Because the numbers of samples associated with the four perceived-realness ratings were not necessarily equal, prediction performance was quantified using balanced accuracy rather than overall accuracy. Balanced accuracy was calculated as the mean of the class-wise accuracies (recalls) across the four perceived-realness categories, yielding a single decoding accuracy for each participant and ROI. The theoretical chance level was 25 %.

#### Perceived realness decoding under the control condition

To examine whether perceived realness could also be decoded from neural responses measured under the control condition, we applied the same decoding procedure described above to the BOLD response patterns acquired in fMRI experiment 1−2. For each stimulus, the perceived-realness rating obtained from the same participant in fMRI experiment 1−1 was used as the target label. Thus, the classifiers predicted the four-level perceived-realness ratings measured in fMRI experiment 1−1 from BOLD activity measured independently in fMRI experiment 1−2. The same source-image-based 10-fold cross-validation procedure, linear SVM classifier, and balanced-accuracy measure as in the main perceived-realness-decoding analysis were used.

#### Perceived realness decoding beyond image features and RT-based decision difficulty

To test whether perceived realness could be decoded from BOLD response patterns beyond variance attributable to image-computable features and decision-related activity, we performed additional residual-decoding analyses. For each stimulus used in fMRI experiment 1−1, we constructed an image-feature vector by concatenating the image statistics described above with compressed VGG-19 style features from layers conv1_1, conv1_2, conv2_1, conv2_2, conv3_1, conv4_1, and conv5_1. Reaction time (RT) for each stimulus presentation was additionally used as a proxy for decision difficulty and decision-related variation.

All residual-decoding analyses used the same source-image-based 10-fold cross-validation scheme as the main perceived-realness-decoding analysis. Thus, all morphs derived from the same original image were assigned to the same outer fold, and stimuli sharing a source-image identity never appeared in both the outer-training and outer-test sets.

To remove BOLD variance predictable from image features, we fitted a ridge-regression encoding model separately for each participant and ROI, predicting the response of each voxel from the concatenated image-feature vector. Image features were standardized using the mean and standard deviation estimated from the corresponding training data only. The ridge regularization parameter was selected independently within each outer-training set using five-fold nested cross-validation, with inner folds also grouped according to source-image identity. Seventeen logarithmically spaced candidate values ranging from 10^−2^ to 10^6^ were evaluated. Model performance was quantified using predictive R^2^, calculated separately for each voxel relative to a baseline prediction given by the mean BOLD response in the corresponding inner-training set and then averaged across voxels. For each candidate regularization parameter, predictive R^2^ was averaged across inner folds. The smallest regularization parameter whose mean predictive R^2^ was within one standard error of the maximum was selected.

Using the selected regularization parameter, an encoding model was fitted to the complete outer-training set and used to predict BOLD responses for the held-out outer-test stimuli. For the outer-training data, out-of-fold BOLD predictions were generated using the inner cross-validation folds, such that the predicted response for each training stimulus was obtained from a model that had not been fitted to that stimulus. Image-feature-residual BOLD patterns were then obtained by subtracting the predicted responses from the observed responses separately for the outer-training and outer-test sets.

In a separate analysis, we removed the BOLD variance related to reaction time (RT). RTs were log-transformed and standardized within each outer cross-validation fold using the mean and standard deviation estimated from the outer-training data. For each voxel, BOLD responses were linearly regressed on standardized log RT using the outer-training data. The fitted model was used to predict the RT-related BOLD component in the outer-test data, which was subtracted from the observed response. RT-related responses in the outer-training set were estimated using five-fold cross-fitting, again grouping the inner folds according to source-image identity, so that the residual response for each training sample was based on a regression model fitted without that sample.

We further constructed a combined residual condition in which image-feature- and RT-related components were removed sequentially. First, the image-feature-predictable component was removed using the ridge-regression procedure described above. The RT-related component was then estimated from the resulting image-feature-residual BOLD responses and removed. For the outer-training data, both residualization steps were performed within the inner cross-validation procedure to generate out-of-fold residuals. For the outer-test data, all model parameters were estimated exclusively from the corresponding outer-training data.

The resulting image-feature-residual, RT-residual, and combined image-feature-plus-RT-residual BOLD patterns were each submitted to the same four-class perceived-realness-decoding procedure described above. Linear SVMs with one-versus-one error-correcting output codes were trained on the residual patterns from the outer-training set and evaluated on the corresponding held-out outer-test set. Predicted labels were pooled across the 10 outer folds, and decoding performance was quantified as balanced accuracy across the four perceived-realness categories.

#### Statistical tests for perceived realness decoding

Statistical significance of perceived-realness-decoding performance was assessed across participants. For each ROI and decoding condition, balanced accuracy was compared with the theoretical chance level of 25 % using a one-sided one-sample *t*-test, with the alternative hypothesis that decoding accuracy was greater than chance. The resulting *p*-values were corrected for multiple comparisons using the Benjamini–Hochberg false-discovery-rate (FDR) procedure^79,80^ across all ROIs and all decoding conditions.

To assess the effect of removing image-feature- and/or RT-related variance from the BOLD responses, decoding accuracy in each residual-decoding condition was compared with that obtained from the original, non-residualized BOLD responses using two-sided paired *t*-tests across participants, separately for each ROI. For these condition comparisons, Benjamini– Hochberg FDR correction was applied jointly across all ROI × condition contrasts. We additionally compared decoding performance across groups of ROIs. ROIs were grouped into early (V1, V2, and V3), ventral (hV4, VO, PPA, and VTC), early dorsal (V3A and V3B), lateral (LO and hMT+), and higher dorsal (V7, IPS1, IPS2, and IPS3−IPS5) visual areas. For each participant, decoding accuracy was first averaged across ROIs within each group. The higher dorsal group was then compared with each of the other ROI groups using two-sided paired *t*-tests. These comparisons were performed separately for each decoding condition, and Benjamini–Hochberg FDR correction was applied across the four ROI-group comparisons within each condition. Statistical significance was determined at an FDR-corrected threshold of *P* < 0.05.

#### Image feature decoding

To examine whether image-computable information could be decoded from BOLD activity, we decoded image features from the BOLD signals recorded in fMRI experiment 2. We separately analysed the image statistics and the style-feature representations extracted from shallow (conv1_1, conv1_2, conv2_1, and conv2_2), middle (conv3_1 and conv4_1), and deep (conv5_1) layers of VGG-19. For each participant and ROI, a regularized linear decoder was trained to predict each dimension of a given image-feature representation from the multivoxel BOLD response pattern. Trials without a valid behavioral response or an available BOLD response pattern were excluded from the analysis. On average, 5.8 % of trials were excluded per participant (range: 0.0–19.1 %).

Decoding was performed using elastic-net regression (α = 0.5). Both the BOLD responses and target feature values were standardized using the means and standard deviations estimated from the training data within each cross-validation fold. A separate decoder was fitted for each feature dimension. The regularization parameter was selected independently for each feature dimension using three-fold inner cross-validation within the training data, with the value minimizing the cross-validated mean squared error selected for the final model. The resulting decoder was then applied to the held-out BOLD responses to reconstruct the corresponding image-feature vectors.

Decoding performance was quantified using pairwise identification accuracy. For each held-out image, the correlation between its predicted feature vector and its true feature vector was compared with the correlation between the same predicted vector and the feature vector of each competing image in the held-out set. A comparison was counted as correct when the predicted feature vector was more strongly correlated with the feature vector of the corresponding image than with that of the competing image; ties were assigned a score of 0.5. Pairwise identification accuracy was calculated as the proportion of correct comparisons and had a theoretical chance level of 50 %.

#### Statistical tests for image feature decoding

Statistical significance of pairwise identification accuracy was assessed across participants using one-sided one-sample *t*-tests against the theoretical chance level of 50 %, with the alternative hypothesis that decoding accuracy was greater than chance. Tests were performed separately for each ROI and image-feature representation using participant-level pairwise identification accuracies. The resulting P-values were jointly corrected for multiple comparisons across all ROIs included in the analysis and all image-feature representations using the Benjamini–Hochberg false-discovery-rate (FDR) procedure. Statistical significance was defined as FDR-corrected *P* < 0.05.

#### Representational similarity analysis

We performed ROI-wise representational similarity analysis (RSA) to examine the correspondence between the representational geometry of BOLD responses and candidate models based on image-computable features, perceived realness, and reaction time (RT). Analyses were conducted separately for each participant and ROI.

To reduce noise in the BOLD response patterns, stimuli were grouped into pseudo-trials according to morph level. For each of the 11 morph levels, the constituent images were randomly divided into two non-overlapping groups of approximately equal size, and multivoxel BOLD response patterns were averaged within each group. This procedure yielded 22 pseudo-trials. To reduce dependence on a particular partition of images, 100 unique pseudo-trial grouping patterns were generated, with the same grouping patterns applied to all participants. Images for which valid BOLD responses, realness ratings, image features, or RTs were unavailable were excluded on a participant-by-participant basis. A pseudo-trial was retained for a participant when at least one valid image remained in that group.

Before pseudo-trial averaging, voxel-wise BOLD responses were standardized across valid images within each participant and ROI. The image-feature model was constructed by concatenating the image statistics and style features described above, and each feature dimension was standardized across images before pseudo-trial averaging. The perceived-realness model was constructed from participant-specific perceived-realness ratings. For the RT model, RTs were log-transformed and standardized within each participant before pseudo-trial averaging.

For each pseudo-trial grouping pattern, a neural representational dissimilarity matrix (RDM) was constructed from the pseudo-trial-averaged multivoxel BOLD patterns using correlation distance (1 − Pearson’s *r*). The image-feature RDM was likewise constructed using correlation distance between pseudo-trial-averaged feature vectors. Because perceived realness and RT were scalar variables, their RDMs were constructed using city-block distance between the corresponding pseudo-trial-averaged values. RDM elements involving unavailable pseudo-trials were treated as missing while retaining a common RDM structure across participants.

For each participant and ROI, the neural RDM was compared separately with each candidate model RDM using Spearman’s rank correlation coefficient. RSA correlations were calculated separately for each of the 100 pseudo-trial grouping patterns and then averaged across grouping patterns to obtain one RSA value for each participant, ROI, and candidate model.

Statistical significance of the RSA correlations was assessed using permutation tests. For each permutation, the pseudo-trial assignments of each candidate model were randomly permuted, the corresponding model RDM was recomputed, and its Spearman correlation with the neural RDM was recalculated. The same permutation procedure was applied to each of the 100 pseudo-trial grouping patterns, and permuted correlations were averaged across grouping patterns in the same manner as the observed correlations. Group-level null distributions were then obtained by averaging the permuted RSA correlations across participants for each permutation. The observed group-mean correlation was tested against the upper tail of the corresponding null distribution using 2,000 permutations. For each candidate model, the resulting *p*-values were corrected for multiple comparisons across the ROIs included in the analysis using the Benjamini–Hochberg false-discovery-rate (FDR) procedure, with statistical significance defined as FDR-corrected *P* < 0.05.

To compare the correspondence of the neural RDM with perceived realness against that with the other candidate models, participant-level RSA correlations were compared within each ROI using two-sided paired-sample *t*-tests. Specifically, the difference between the perceived-realness RSA correlation and each other model RSA correlation was tested against zero across participants. The resulting *p*-values were jointly corrected using the Benjamini–Hochberg FDR procedure across all ROIs included in the analysis and all perceived-realness-versus-other-model comparisons, with FDR-corrected *P* < 0.05 considered significant.

#### Linear trend of model comparison

To test whether the relative correspondence with perceived realness increased along the dorsal visual hierarchy, we conducted a linear-trend analysis on the difference between the RSA correlations for the perceived-realness and image-feature models. For each participant and ROI, the difference score was calculated by subtracting the Spearman correlation for the image-feature model from that for the perceived-realness model.

ROIs were grouped into three anatomically defined stages: early visual areas (V1, V2, and V3), early dorsal visual areas (V3A and V3B), and higher dorsal visual areas (V7, IPS1, IPS2, and IPS3−IPS5). For each participant, difference scores were averaged across ROIs within each stage. A linear regression was then fitted to the three stage-level difference scores using equally spaced ordinal positions (1, 2, and 3), yielding a participant-specific slope. Only participants with valid RSA estimates for all ROIs included in the trend analysis were entered into the statistical test. The resulting slopes were tested against zero using a one-sided one-sample *t*-test, with the alternative hypothesis that the mean slope was greater than zero, corresponding to an increase in the relative correspondence with the perceived-realness model from early visual to higher dorsal areas. Statistical significance was determined at *P* < 0.05.

#### Bootstrap-based model winner probability

To characterize the relative ranking of the candidate RSA models, we performed a participant-level bootstrap analysis separately for each ROI. For each participant, ROI, and model, we used the Spearman RSA correlation averaged across the 100 pseudo-trial grouping patterns. In each of 2,000 bootstrap iterations, participants were sampled with replacement, with the bootstrap sample size equal to the number of participants included in the analysis. For each candidate model, the mean RSA correlation was calculated across the resampled participants, and the model with the highest mean correlation was designated as the winner for that iteration. The winner probability for each model was defined as the proportion of bootstrap iterations in which that model achieved the highest mean RSA correlation.

The same analysis was also performed for anatomically defined ROI groups. For each participant and candidate model, RSA correlations were first averaged across the ROIs belonging to each ROI group, after which participants were resampled with replacement and model winner probabilities were calculated using the same procedure. Winner probabilities were treated as descriptive measures of the stability of model ranking under participant resampling and were not subjected to significance testing.

### MEG acquisition and preprocessing

#### MEG acquisition

MEG data were acquired in a magnetically shielded room at the NICT CiNet imaging facility (Suita, Osaka, Japan) using a customized 360-channel whole-head MEG system (Neuromag 360, MEGIN-Elekta). The system consisted of 204 planar gradiometers, 102 magnetometers, and 54 axial gradiometers. MEG signals were recorded at a sampling rate of 1,000 Hz. The 204 planar gradiometers were arranged as orthogonal pairs at 102 sensor locations, measuring magnetic-field gradients in two orthogonal directions. Head position relative to the sensor array was determined using five head-position indicator coils attached to the forehead and preauricular regions. The positions of the coils were measured using a three-dimensional digitizer. Electrooculograms (EOGs) were recorded using four electrodes placed approximately 1.5 cm above and below the left eye and at the left and right temples. The EOG signals were recorded simultaneously with the MEG signals through auxiliary input channels of the MEG system. Bad MEG channels were automatically detected and rejected using the internal algorithm implemented in the MaxFilter package. EOG signals were additionally used to visually inspect the data for artefacts related to eye movements. The timing of MEG acquisition was synchronized with stimulus presentation on a trial-by-trial basis using a photosensor placed in the upper-right corner of the display over a white square that appeared concurrently with stimulus presentation.

#### Preprocessing

MEG data were band-pass filtered between 0.1 and 45 Hz. Independent component analysis (ICA) was subsequently applied, and components associated with eye movements were identified with reference to the simultaneously recorded EOG signals and removed from the data.

#### MEG source reconstruction

Source reconstruction was performed using MNE-Python (https://mne.tools/stable/index.html; Ref. 81). For each participant, an individual cortical source space with octahedral spacing (oct6) and an individual boundary element model (BEM) were used to compute the forward solution. All available MEG channels in the preprocessed data were used for source reconstruction. A minimum-norm inverse operator was constructed using a loose orientation constraint of 0.2 and depth weighting of 0.8. For the stimulus-locked analysis, epochs were extracted from 200 ms before to 1,000 ms after stimulus onset, with stimulus onset determined from the photodiode signal. Epochs were baseline-corrected using the prestimulus interval from −200 to 0 ms, and the noise covariance matrix was estimated empirically from the same prestimulus interval. Within each run and experimental condition, trials were divided into approximately equally sized bundles of approximately 15–20 trials, and sensor-level MEG signals were averaged across trials within each bundle before source reconstruction. Source activity was estimated separately for each averaged response using dynamic statistical parametric mapping (dSPM), with a regularization parameter of λ^2^ = 1/9, corresponding to an assumed signal-to-noise ratio of 3. Vector source estimates were computed at each cortical location, and the magnitude across the three orientation components was used for subsequent analyses. For each individually defined ROI, source estimates from all vertices contained within the ROI were retained without additional activity-based vertex selection. For the response-locked analysis, response events were identified from the recorded button-response channels and matched to the corresponding stimulus presentations. When multiple responses occurred within a trial, only the first response following stimulus onset was retained. Trials without a recorded response were not included in the response-locked analysis. Epochs were extracted from 1,000 ms before to 500 ms after the response. No baseline correction was applied to the response-locked epochs. Instead, the noise covariance matrix was estimated empirically from the −200 to 0 ms interval preceding stimulus onset, thereby using the same prestimulus period as in the stimulus-locked analysis. Trials were grouped and averaged within each run and response condition using the same procedure as for the stimulus-locked analysis, and source reconstruction was performed using the same forward model, inverse parameters, and dSPM procedure described above.

### MEG data analysis

#### Perceived realness (real or unreal) decoding

To examine the temporal dynamics of neural information distinguishing real from unreal images, we performed time-resolved binary decoding separately for each participant using stimulus-locked, source-localized MEG activity from V1, V3A, and IPS3−IPS5. For each ROI, source activity from all vertices in the left and right hemispheres was concatenated along the vertex dimension, without averaging across hemispheres. Decoding was restricted to source activity from 1 to 500 ms after stimulus onset.

Decoding was performed within 20-ms sliding time windows advanced in 10-ms steps. Within each window, activity from all vertices in both hemispheres and all time points within the window was concatenated to form a single feature vector for each sample. To reduce dimensionality, principal component analysis (PCA) was fitted using the training data only within each cross-validation fold. The smallest number of principal components accounting for at least 99 % of the cumulative variance in the training data was retained. The held-out data were projected onto the same components using the mean and component loadings estimated exclusively from the training data.

Real versus unreal images were classified using a linear support vector machine implemented with the MATLAB function ‘fitcsvm’. To prevent MEG responses to the same image from appearing in both the training and test sets, leave-one-run-out cross-validation was performed separately within each experimental day (8 runs per day). In each fold, one run was held out for testing, and the remaining seven runs from the same day were used for training. Decoding performance in each fold was quantified as balanced accuracy, calculated as the mean recall across the real and unreal classes. Balanced accuracies were then averaged across all folds and both days to obtain a single participant-level decoding accuracy for each ROI and time window. The theoretical chance level was 50 %. Statistical significance of decoding performance was assessed across participants separately for each ROI and time window using a one-sided one-sample *t*-test against the 50 % chance level, with the alternative hypothesis that decoding accuracy was greater than chance. The resulting p-values were corrected for multiple comparisons using the Benjamini–Hochberg false-discovery-rate (FDR) procedure jointly across all ROIs and all time windows. Statistical significance was determined at an FDR-corrected threshold of *P* < 0.05.

#### Statistical tests for latency separation across ROIs

To compare the timing of peak perceived-realness decoding across visual areas, peak latencies were estimated using a participant-bootstrap procedure^10,21^. Analyses focused on V1, V3A, and IPS3–IPS5. For each ROI, peak latency was defined as the time point at which the group-average decoding accuracy reached its maximum within the stimulus-locked analysis interval. No additional temporal smoothing was applied for the peak-latency analysis. To estimate uncertainty in individual peak latencies, participants were resampled with replacement 10,000 times, with each bootstrap sample containing the same number of participants as the original sample. For each bootstrap sample, the group-average decoding time course was recomputed and its peak latency was identified. The 2.5th and 97.5th percentiles of the resulting bootstrap distribution were used to define the 95 % confidence interval of the peak latency.

To test whether perceived-realness decoding peaked later in higher dorsal visual areas, peak-latency differences were assessed between IPS3–IPS5 and V1 and between IPS3– IPS5 and V3A. Within each bootstrap iteration, the same resampled set of participants was used for all ROIs, thereby preserving the paired structure of the data. The peak-latency difference was calculated as the peak latency in IPS3–IPS5 minus that in the comparison ROI. For each comparison, a one-sided bootstrap test assessed the directional hypothesis that decoding peaked later in IPS3–IPS5 than in the comparison ROI. The P value was calculated as the proportion of bootstrap samples in which the peak-latency difference was less than or equal to zero. The 95 % confidence interval of each peak-latency difference was defined by the 2.5th and 97.5th percentiles of its bootstrap distribution. The resulting P values for the two planned comparisons were corrected for multiple comparisons using the Benjamini–Hochberg false-discovery-rate (FDR) procedure, with FDR-corrected *P* < 0.05 considered significant.

#### Image feature decoding from MEG source activity

To examine the temporal dynamics of image-computable information in cortical activity, we decoded image-feature representations from stimulus-locked, source-localized MEG signals. The image statistics and shallow, middle, and deep style-feature representations were analysed separately. For each MEG sample, the target feature vector was obtained by averaging the corresponding feature vectors across the images included in that sample.

Decoding was restricted to source activity from 1 to 500 ms after stimulus onset. Decoding was performed separately for each participant, ROI, feature representation, and 20-ms time window, with consecutive windows advanced in 10-ms steps. Within each time window, source activity from all vertices and time points was concatenated to form the predictor vector. To prevent MEG responses to the same images from appearing in both the training and test sets, leave-one-run-out cross-validation was performed separately within each experimental day (8 runs per day). In each fold, one run was held out for testing, and the remaining seven runs from the same day were used for training. Prediction performance was averaged across all folds and both days to obtain a single participant-level decoding score for each ROI, feature representation, and time window. To reduce the dimensionality of the MEG patterns, principal component analysis (PCA) was fitted using only the outer-training data in each cross-validation fold. The smallest number of principal components accounting for at least 99 % of the cumulative variance in the training data was retained. The held-out data were projected onto the retained components using the mean and component loadings estimated exclusively from the corresponding training data.

A separate elastic-net regression model (α = 0.5) was fitted to predict each dimension of the image-feature representation from the MEG source pattern. Both the MEG predictors and target feature values were standardized using parameters estimated exclusively from the outer-training data. A common regularization parameter was used across feature dimensions within each participant, ROI, time window, and outer cross-validation fold. This parameter was determined from five representative feature dimensions sampled across the target feature vector. For each representative dimension, eight candidate regularization values were evaluated using three-fold cross-validation within the outer-training data, and the value minimizing cross-validated mean squared error was selected. The common regularization parameter was then defined as the geometric mean of the selected values across the valid representative feature dimensions and was used to fit all feature-specific decoders.

Decoding performance was quantified using pairwise identification accuracy within each held-out run. For each predicted feature vector, its Pearson correlation with the feature vector of the corresponding sample was compared with its correlation with the feature vector of each other sample in the same held-out run. A comparison was scored as correct when the correlation with the corresponding sample was higher, incorrect when it was lower, and 0.5 in the case of a tie. Pairwise identification accuracy was calculated as the mean of these scores and had a theoretical chance level of 50 %. Participant-level accuracy for each ROI, time window, and feature representation was obtained by averaging pairwise identification accuracy across the leave-one-run-out folds.

Statistical significance of pairwise identification accuracy was assessed across participants using one-sided one-sample *t*-tests against the 50 % chance level, with the alternative hypothesis that decoding accuracy was greater than chance. Multiple comparisons were controlled using the Benjamini–Hochberg false-discovery-rate (FDR) procedure. Correction was performed separately for each ROI, with all time windows and all four image-feature representations included jointly in a single correction family for that ROI. Statistical significance was determined at an FDR-corrected threshold of *P* < 0.05. For summaries of decoding onset, sustained significant decoding was defined as beginning at the first of at least two consecutive time windows reaching the FDR-corrected significance threshold.

#### Response-locked decoding and peak-timing analysis

To examine the temporal relationship between perceived-realness-related neural information and behavioral responses, we additionally performed time-resolved real-versus-unreal decoding using response-locked source-localized MEG activity. The decoding procedure was otherwise identical to that used for the stimulus-locked analysis. Decoding was restricted to source activity from -999 to 500 ms relative to response onset. Decoding was performed separately for each participant and ROI using 20-ms sliding time windows advanced in 10-ms steps. Within each time window, source activity from all vertices and time points was concatenated to form the feature vector for each sample.

Prediction performance was evaluated using leave-one-run-out cross-validation separately within each experimental day (8 runs per day). Within each cross-validation fold, principal component analysis (PCA) was fitted exclusively to the training data, and the smallest number of principal components accounting for at least 99 % of the cumulative variance was retained. The held-out data were projected using the mean and component loadings estimated from the corresponding training data. Real and unreal samples were then classified using a linear support vector machine, and performance was quantified using balanced accuracy, defined as the mean recall across the two classes. Participant-level decoding accuracy was obtained by averaging balanced accuracy across cross-validation folds, with a theoretical chance level of 50 %.

Statistical significance of response-locked decoding performance was assessed across participants using one-sided one-sample *t*-tests against the 50 % chance level, with the alternative hypothesis that decoding accuracy was greater than chance. Because statistical testing of response-locked decoding was restricted to IPS3–IPS5 and to time windows whose centers fell between −500 and 500 ms relative to response onset, the resulting *P*-values were corrected for multiple comparisons across all tested response-locked time windows using the Benjamini–Hochberg false-discovery-rate (FDR) procedure. Statistical significance was determined at an FDR-corrected threshold of *P* < 0.05.

To determine whether the response-locked decoding peak in IPS3–IPS5 occurred after response onset, peak latency was estimated using a participant-bootstrap procedure10,21. Peak estimation was restricted to the interval from −500 ms to the end of the response-locked analysis window relative to response onset. Peak latency was defined as the time point at which the group-average decoding accuracy reached its maximum within this interval. No additional temporal smoothing or fractional-peak estimation was applied. To estimate uncertainty in the group-level peak latency, participants were resampled with replacement 10,000 times, with each bootstrap sample containing the same number of participants as the original sample. For each bootstrap sample, the group-average response-locked decoding time course was recalculated and its peak latency was identified using the same procedure. The 2.5th and 97.5th percentiles of the resulting bootstrap distribution defined the 95 % confidence interval of the peak latency.

The directional hypothesis that the response-locked decoding peak occurred after response onset was tested using the same bootstrap distribution. Specifically, the one-sided P-value was calculated as the proportion of bootstrap samples in which the estimated peak latency was less than or equal to 0 ms, corresponding to the null boundary of response onset. Statistical significance was determined at *P* < 0.05.

## Author contributions

T.O. and H.B. conceived the study. T.O. conceived the original idea. T.O. and H.B. designed the experiments and developed the methodology. T.O. conducted the experiments, collected and curated the data, and performed the formal analyses. T.O. and H.B. acquired funding. H.B. provided resources and supervised the project. T.O. wrote the original draft of the manuscript. Both authors reviewed and edited the manuscript.

## Acknowledgements

We thank Dr. Roland W. Fleming and Fleming Lab members for discussions and suggestions. We also thank Ms. Naomi Murakami and Ms. Miho Hayashi for their assistance with the fMRI and MEG experiments, and Mr. Ichiro Fujimoto, Dr. Hironori Nishimoto, Dr. Asuka Otsuka, Dr. Tomoki Haji, Mr. Takanori Mimura, Dr. Takeshi Nogai, Mr. Koji Uchida, Mr. Daisuke Nishiyama, and Ms. Yuriko Nohara for the operations of fMRI and MEG. This study is supported by JSPS KAKENHI 24KJ2222, 25K17200, and 25KK0044 (to T.O.), and JSPS KAKENHI 21H00968 and JST ERATO JPMJER1801 (to H.B.).

## Data Availability

The anonymized data supporting the findings of this study will be made publicly available through an appropriate repository upon publication of the peer-reviewed article.

## Code Availability

The analysis code supporting the findings of this study will be made publicly available through an appropriate repository upon publication of the peer-reviewed article.

## Declaration on the use of generative AI tools for writing

During the preparation of this manuscript, the authors used ChatGPT (OpenAI; GPT-5.5 and GPT-5.6) to assist with translating passages initially drafted in their native language, correcting grammar, and improving the clarity and readability of the English text. All AI-assisted text was carefully reviewed and substantively revised by the authors before inclusion in the manuscript. The tool was not used to generate scientific ideas, arguments, interpretations, or conclusions. The authors take full responsibility for the accuracy, integrity, and content of the manuscript.

## Supporting information

### Details of fMRI preprocessing

In accordance with the reporting guidelines for fMRIPrep, we provide below the methods description automatically generated by fMRIPrep.

## Anatomical data preprocessing

A total of 1 T1-weighted (T1w) images were found within the input BIDS dataset. The T1w image was corrected for intensity non-uniformity (INU) with N4BiasFieldCorrection (Tustison et al. 2010), distributed with ANTs 2.5.0 (Avants et al. 2008, RRID:SCR_004757), and used as T1w-reference throughout the workflow. The T1w-reference was then skull-stripped with a Nipype implementation of the antsBrainExtraction.sh workflow (from ANTs), using OASIS30ANTs as target template. Brain tissue segmentation of cerebrospinal fluid (CSF), white-matter (WM) and gray-matter (GM) was performed on the brain-extracted T1w using fast (FSL (version unknown), RRID:SCR_002823, Zhang, Brady, and Smith 2001). Brain surfaces were reconstructed using recon-all (FreeSurfer 7.3.2, RRID:SCR_001847, Dale, Fischl, and Sereno 1999), and the brain mask estimated previously was refined with a custom variation of the method to reconcile ANTs-derived and FreeSurfer-derived segmentations of the cortical gray-matter of Mindboggle (RRID:SCR_002438, Klein et al. 2017). Volume-based spatial normalization to one standard space (MNI152NLin2009cAsym) was performed through nonlinear registration with antsRegistration (ANTs 2.5.0), using brain-extracted versions of both T1w reference and the T1w template. The following template was selected for spatial normalization and accessed with TemplateFlow (23.1.0, Ciric et al. 2022): ICBM 152 Nonlinear Asymmetrical template version 2009c [Fonov et al. (2009), RRID:SCR_008796; TemplateFlow ID: MNI152NLin2009cAsym].

## Functional data preprocessing

For each run found per subject (across all tasks and sessions), the following preprocessing was performed. First, a reference volume was generated, using a custom methodology of fMRIPrep, for use in head motion correction. Head-motion parameters with respect to the BOLD reference (transformation matrices, and six corresponding rotation and translation parameters) are estimated before any spatiotemporal filtering using mcflirt (FSL , Jenkinson et al. 2002). The BOLD reference was then co-registered to the T1w reference using bbregister (FreeSurfer) which implements boundary-based registration (Greve and Fischl 2009). Co-registration was configured with six degrees of freedom. Several confounding time-series were calculated based on the preprocessed BOLD: framewise displacement (FD), DVARS and three region-wise global signals. FD was computed using two formulations following Power (absolute sum of relative motions, Power et al. (2014)) and Jenkinson (relative root mean square displacement between affines, Jenkinson et al. (2002)). FD and DVARS are calculated for each functional run, both using their implementations in Nipype (following the definitions by Power et al. 2014). The three global signals are extracted within the CSF, the WM, and the whole-brain masks. Additionally, a set of physiological regressors were extracted to allow for component-based noise correction (CompCor, Behzadi et al. 2007). Principal components are estimated after high-pass filtering the preprocessed BOLD time-series (using a discrete cosine filter with 128s cut-off) for the two CompCor variants: temporal (tCompCor) and anatomical (aCompCor). tCompCor components are then calculated from the top 2% variable voxels within the brain mask. For aCompCor, three probabilistic masks (CSF, WM and combined CSF+WM) are generated in anatomical space. The implementation differs from that of Behzadi et al. in that instead of eroding the masks by 2 pixels on BOLD space, a mask of pixels that likely contain a volume fraction of GM is subtracted from the aCompCor masks.

This mask is obtained by dilating a GM mask extracted from the FreeSurfer’s aseg segmentation, and it ensures components are not extracted from voxels containing a minimal fraction of GM. Finally, these masks are resampled into BOLD space and binarized by thresholding at 0.99 (as in the original implementation). Components are also calculated separately within the WM and CSF masks. For each CompCor decomposition, the k components with the largest singular values are retained, such that the retained components’ time series are sufficient to explain 50 percent of variance across the nuisance mask (CSF, WM, combined, or temporal). The remaining components are dropped from consideration. The head-motion estimates calculated in the correction step were also placed within the corresponding confounds file. The confound time series derived from head motion estimates and global signals were expanded with the inclusion of temporal derivatives and quadratic terms for each (Satterthwaite et al. 2013). Frames that exceeded a threshold of 0.5 mm FD or 1.5 standardized DVARS were annotated as motion outliers. Additional nuisance timeseries are calculated by means of principal components analysis of the signal found within a thin band (crown) of voxels around the edge of the brain, as proposed by (Patriat, Reynolds, and Birn 2017). All resamplings can be performed with a single interpolation step by composing all the pertinent transformations (i.e. head-motion transform matrices, susceptibility distortion correction when available, and co-registrations to anatomical and output spaces). Gridded (volumetric) resamplings were performed using nitransforms, configured with cubic B-spline interpolation.

---

**Supplementary Figure 1.**
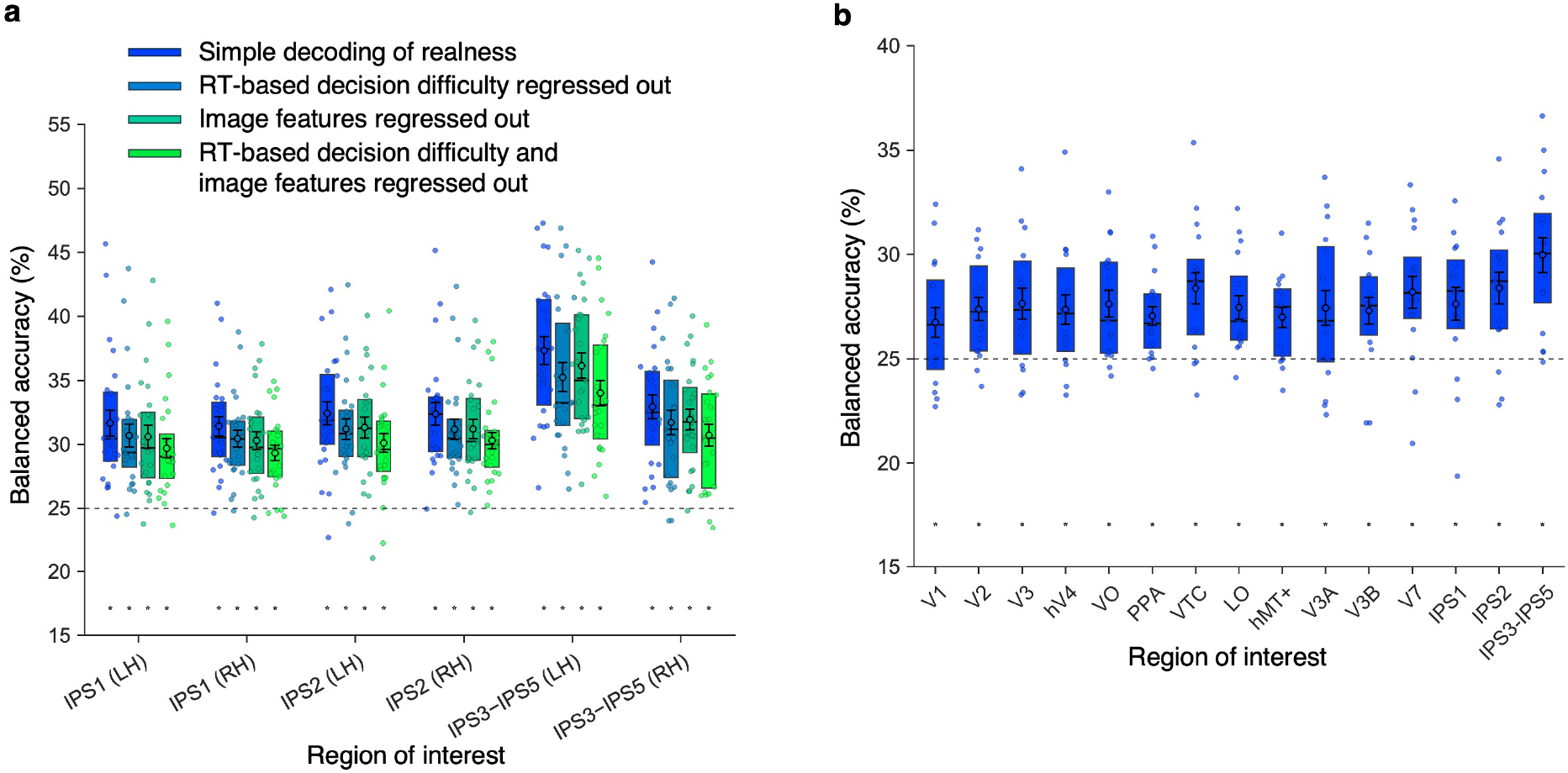
fMRI decoding results in control conditions. **(a)** Decoding results for each hemisphere of IPS1, IPS2, and IPS3–IPS5. Plotting conventions and statistical annotations are the same as in Fig. 3b. **(b)** Decoding results from fMRI experiment 1−2. Asterisks indicate statistical significance using the same convention as in Fig. 3b.

**Supplementary Figure 2.**
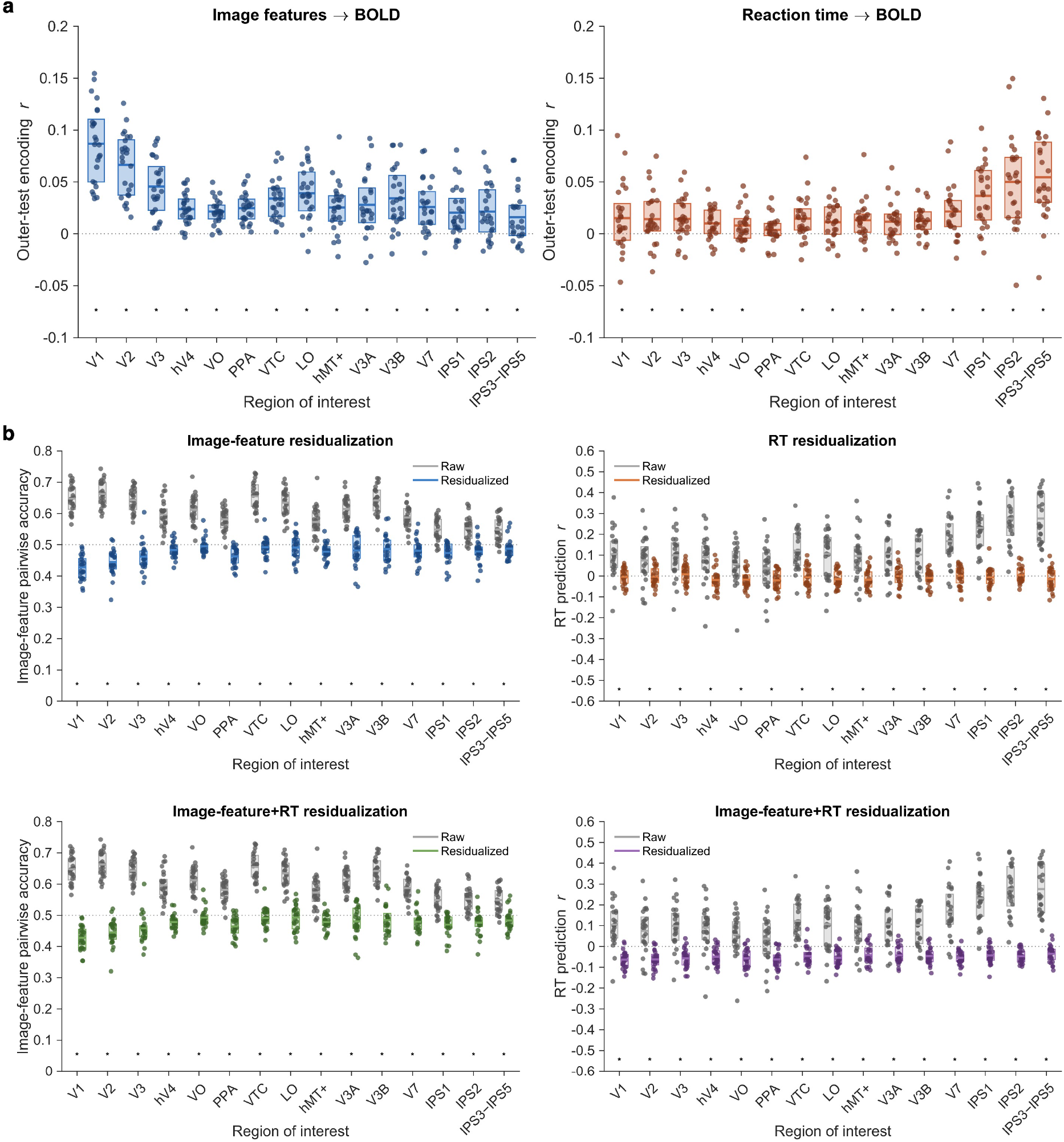
Validation of encoding models. **(a)** Model fit on held-out test data. The left panel shows the fit of the image-feature encoding models, and the right panel shows the fit of the RT-based encoding model. **(b)** Reduction in prediction accuracy for image features and RT after residualization. The upper left panel shows the results after removing image-feature-related components, the upper right panel shows the results after removing the RT-related component, and the two lower panels show the results after removing both components (image-feature prediction on the left and RT prediction on the right).

**Supplementary Figure 3.**
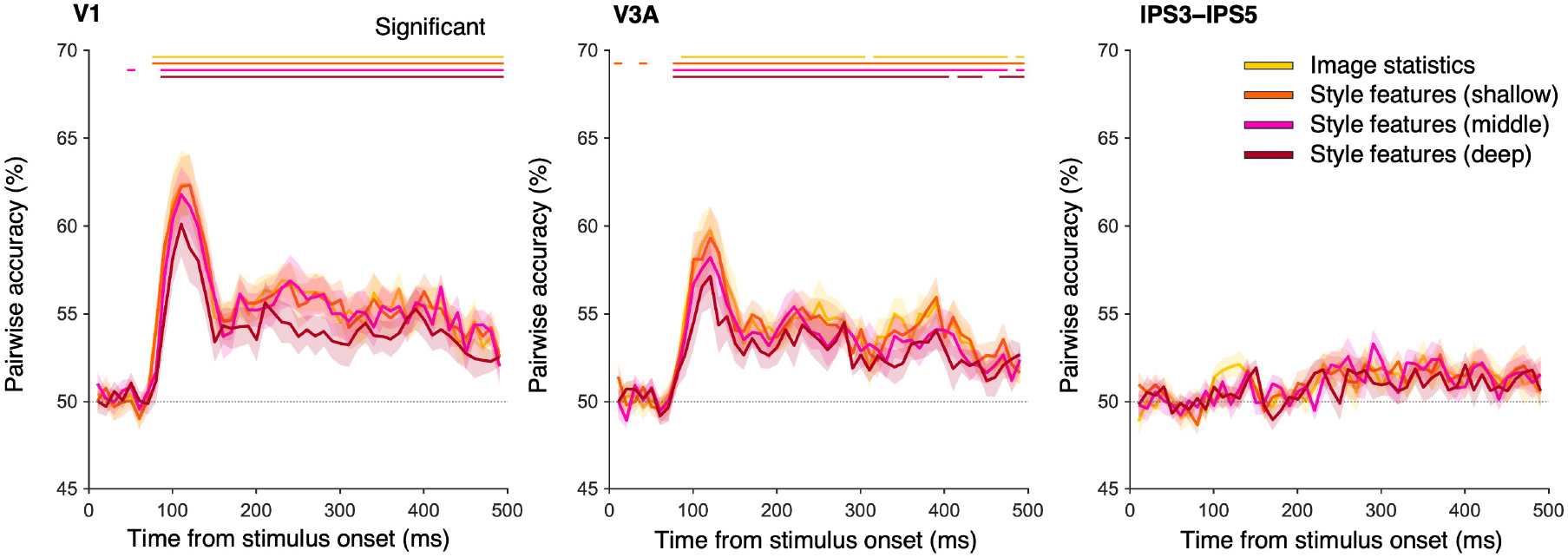
MEG decoding of image features. Decoding results for each class of image features from stimulus-locked, source-localized MEG signals in V1, V3A, and IPS3–IPS5. For each ROI, the yellow, orange, pink, and dark red lines show pairwise accuracy for image statistics and style features extracted from shallow, middle, and deep layers, respectively. Bars above the plots indicate time points at which decoding accuracy was significantly above the theoretical chance level for pairwise accuracy (50 %; FDR-adjusted *P* < 0.05).

